# Exercise reprograms the inflammatory landscape of multiple stem cell compartments during mammalian aging

**DOI:** 10.1101/2022.01.12.475145

**Authors:** Ling Liu, Soochi Kim, Matthew T. Buckley, Jaime M. Reyes, Jengmin Kang, Lei Tian, Mingqiang Wang, Alexander Lieu, Michelle Mao, Cristina Rodriguez-Mateo, Heather Ishak, Mira Jeong, Joseph C. Wu, Margaret A. Goodell, Anne Brunet, Thomas A. Rando

## Abstract

Exercise has the ability to rejuvenate stem cells and improve tissue homeostasis and regeneration in aging animals. However, the cellular and molecular changes elicited by exercise have not been systematically studied across a broad range of cell types in stem cell compartments. To gain better insight into the mechanisms by which exercise affects niche and stem cell function, we subjected young and old mice to aerobic exercise and generated a single cell transcriptomic atlas of muscle, neural and hematopoietic stem cells with their niche cells and progeny.

Complementarily, we also performed whole transcriptome analysis of single myofibers from these animals. We identified common and unique pathways that are compromised across these tissues and cell types in aged animals. We found that exercise has a rejuvenating effect on subsets of stem cells, and a profound impact in the composition and transcriptomic landscape of both circulating and tissue resident immune cells. Exercise ameliorated the upregulation of a number of inflammatory pathways as well as restored aspects of cell-cell communication within these stem cell compartments. Our study provides a comprehensive view of the coordinated responses of multiple aged stem cells and niche cells to exercise at the transcriptomic level.

## INTRODUCTION

Aging leads to stem cell dysfunction resulting in deterioration in tissue homeostasis and regeneration (Brunet and Rando, 2017; Chandel et al., 2016; Goodell and Rando, 2015; Rando and Wyss-Coray, 2014; Ren et al., 2017). Stem cell aging is manifested distinctly at the cellular and molecular levels in different tissues. For example, with age, neural stem cells (NSCs) are depleted in the brain hippocampus and subventricular zone (SVZ), hematopoietic stem cells (HSCs) skew towards the myeloid lineage, and skeletal muscle stem cells (MuSCs) exhibit increased susceptibility to cell death and delayed cell activation in response to injury (Gage and Temple, 2013; Jones and Rando, 2011; Liu and Rando, 2011; Navarro Negredo et al., 2020). In addition, aging elicits distinct alterations across the different cell types in a stem cell niche. Increased heterogeneity in gene expression has been found in various stem cells and the niche cells (Hernando-Herraez et al., 2019; Mahmoudi et al., 2019; Salzer et al., 2018; Xie et al., 2020). To delineate the complexity of aging at the stem cell, tissue and organismal level, it is critical to examine changes in the regulatory molecules and pathways simultaneously across cell types and tissues in aging animals.

Physical exercise has the health benefit of antagonizing the metabolic and immune changes that underlie various age-related diseases, lowering the mortality risk from cardiovascular diseases and cancer (Saint-Maurice et al., 2019). In patients with type II diabetics, physical exercise leads to significant reduction in the plasma Hemoglobin A_1c_ level, an indication of improved glycemic control (Sigal et al., 2007). In people with dyslipidemia, the amount of physical activity positively correlates with the degree of improvement in their lipoprotein profile (Kraus et al., 2002). Even in people with symptomatic but stable coronary artery disease, an increase in regular physical activity leads to increased expression of total endothelial nitric oxide synthase (eNOS), enhanced vasodilatory responses to acetylcholine, and regression of atherosclerotic lesions (Hambrecht et al., 2003). Immunosenescence is a feature of aging and related to an increased susceptibility to infections, autoimmune diseases, chronic inflammation, cancer, and neurologic disorders. Recent evidence suggests that physical exercise has the ability to remodel the immune system during aging and therefore shape the immune landscape in later life (Nieman and Wentz, 2019). In addition, physical activity is associated with lower risk of dementia and neurodegenerative diseases including Alzheimer’s disease and Parkinson’s disease (Fang et al., 2018; Rovio et al., 2005; Scarmeas et al., 2009).

Increasing evidence indicates that physical exercise also has beneficial effects on stem cell function, tissue homeostasis and regeneration in adult animals. Running enhances neurogenesis and learning in the adult mouse brain, and counters the reduction of hippocampal neurogenesis during aging (De Miguel et al., 2021; Horowitz et al., 2020; van Praag et al., 2005). MuSCs in aged animals exhibit reduced expression of Cyclin D1 and a delay in activation in response to acute muscle injury (Brett et al., 2020). Exercise increases Cyclin D1 expression and enhances muscle regeneration in old animals (Brett *et al*., 2020). A recent study identifies a subset of osteogenic progenitors in the bone marrow as niche cells for the common lymphoid progenitors (CLPs) which diminish with age (Shen et al., 2021). Voluntary running increased the number of both the osteogenic progenitors and CLPs in aging animals without affecting the frequency of the HSC population (Shen *et al*., 2021). However, up to date, studies on the mechanisms by which exercise modulates stem cell function have generally not included effects of exercise on local niche cells or endocrine influence from distal cells that are not in direct contact with stem cells. It remains unclear how niche cells of various stem cell compartment are affected in ways that can exert a combinatorial influence on stem cell function following exercise. It is also largely unknown how changes induced by exercise in skeletal muscle, the direct target and beneficiary of physical exercise, are mechanistically connected to changes in distal tissues and the overall health of an individual. A comprehensive examination of the molecular changes induced by exercise across tissue and cell types is critical to understand the systemic effect of exercise in reshaping the molecular and cellular processes in stem cells and their niche environment to improve tissue homeostasis and regeneration during aging.

The development of single cell RNA sequencing (scRNA-seq) technology has led to the creation of atlases of tissues, including those from aged animals, which report changes in the transcriptome of individual cells across multiple tissues and organs (Martinez-Jimenez et al., 2017; Tabula Muris, 2020). The effect of caloric restriction on the cellular composition and transcriptional network of aged tissues has also been examined at the single cell level (Ma et al., 2020). It remains unknown whether other forms of aging intervention exert similar systemic effect across tissues as caloric restriction. Changes in the interaction between stem cells and their niche during aging and in response to aging interventions have not been explored. In this study, we generated an integrative single cell atlas to understand the effect of exercise in modulating stem cell function and tissue homeostasis during aging. We systematically analyzed 435,628 single cells from skeletal muscle, the SVZ of the brain, bone marrow HSPCs and peripheral blood immune cells of young and old mice with and without a period of voluntary running exercise. We analyzed cell type composition changes, transcription changes and intercellular communication changes in these stem cell compartments. Although exercise induced a much smaller subset of genes in aged animals, it was able to ameliorate the elevated inflammation in cell types from all these stem cell compartments. Computational analysis of predicted ligand-receptor interaction also suggests that intercellular communication can be restored by exercise.

In addition, we have performed RNA-seq to understand gene expression changes in single muscle fibers, the contractile units that not only generate the movements that are the basis of exercise-mediated effects, but are also the key niche cell type of MuSCs. We find that exercise largely erases the age-difference in the transcriptome of muscle fibers. Overall, our study provides a comprehensive molecular framework for the effect of exercise in modulating cellular and molecular processes during aging.

## RESULTS

### Construction of an aging and exercise mouse single-cell atlas by scRNA-seq

To understand the transcriptional changes induced by aerobic exercise in stem cells and their associated niche cells in muscular, neural, and hematopoietic systems of young and aged animals, young (“Y”; 4 months of age) and old (“O”; 22 months of age) male C57BL/6J mice were singly housed in cages in which rotating running wheels were attached to digital recorders (Figure 1A). Age-matched control mice were housed in identical cages in which the wheels were removed. All mice provided with the running wheels adapted to a voluntary exercise routine during the dark phase of the sleep-wake cycle within one week of exposure. We harvested skeletal muscle, the subventricular zone (SVZ) of brain, bone marrow, and blood from control mice and from these exercised mice after 5 weeks of running. We processed the tissues into single-cell suspensions and eliminated dead cells by FACS. Due to the rarity of HSCs in the bone marrow, we used a cocktail of antibodies against surface markers of hematopoietic stem and progenitor cells (HSPCs) to enrich for this population. Live single cells were then captured and a total of 458,754 cells from the 4 tissues were sequenced. After quality control, we retained 435,628 cells in our single-cell atlas for downstream analysis.

**Figure 1:**
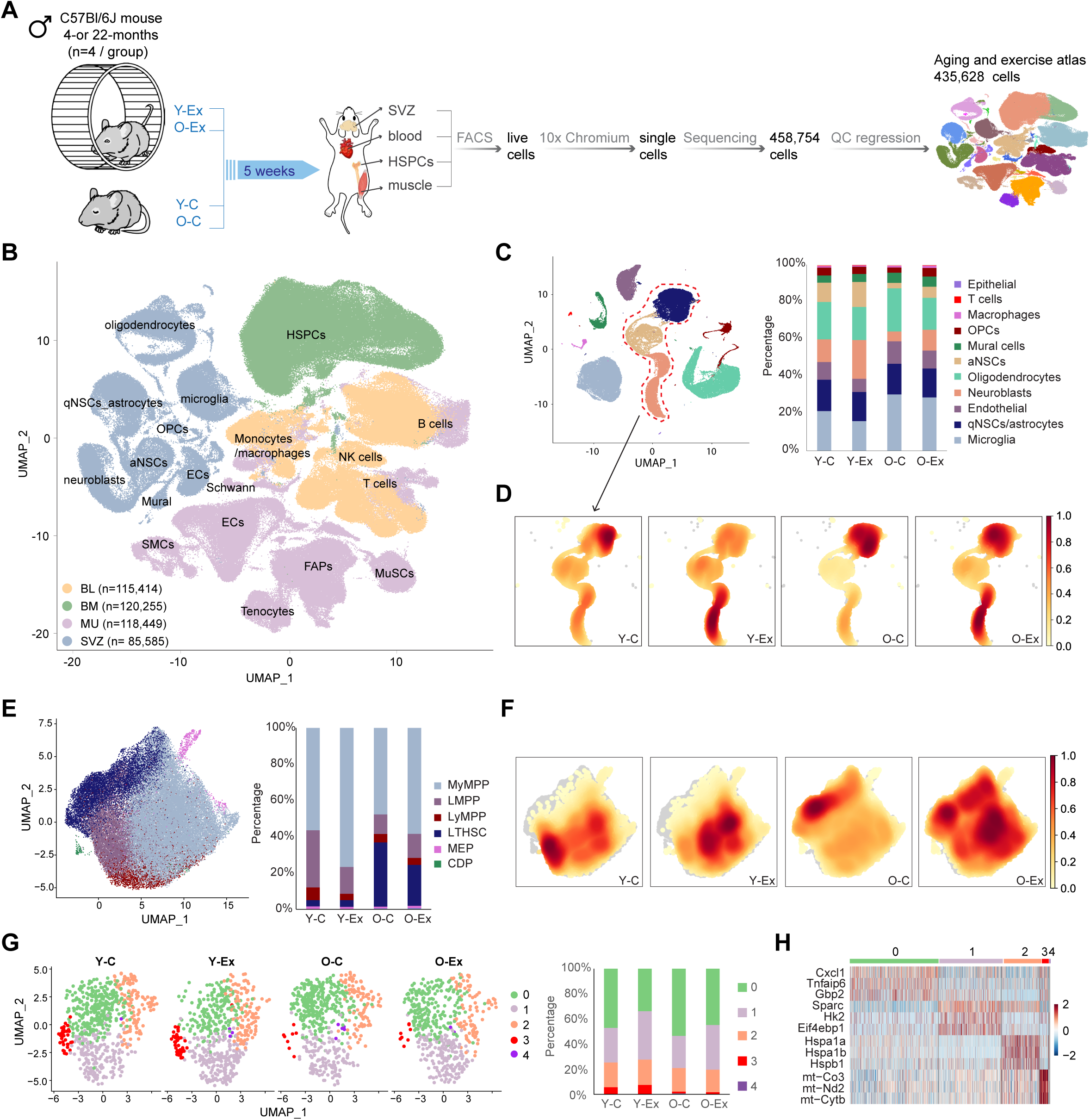
Changes in three stem cell compartments in response to aging and exercise revealed by scRNA-seq. (**A**) Schematic diagram of the multi-tissue scRNA-seq experimental design. Y-C: young control mice; O-C: old control mice; Y-Ex: young exercised mice; O-Ex: old exercised mice; SVZ: subventricular zone; HSPCs: hematopoietic stem and progenitor cells, FACS: fluorescence-activated cell sorting. (**B**) UMAP of the multi-tissue exercise single cell atlas. Cell clusters are colored by the tissue type. The total number of cells in each tissue is indicated in the legend. BL: blood; BM: bone marrow; MU: muscle; ECs: endothelial cells; FAPs: fibro-adipogenic progenitors; OPCs: oligodendrocyte progenitor cells; MuSCs: muscle stem cells; aNSCs: activated neural stem cells; qNSCs: quiescent neural stem cells; SMCs: smooth muscle cells. (**C**) UMAP of cell types in the SVZ. Fraction of each cell type is shown on the right. Neural stem and progenitor cell clusters are circled and used in the density calculation presented in (D). (**D**) Cell density plots of the qNSCs, aNSCs and neuroblasts in control or exercised young and old mice. The color scale represents cell density normalized to a scale of 0 to 1 where 0 indicates an absence of cells and 1 represents the highest cell density. (**E**) UMAP of the HSPCs. Fraction of each cell type is shown on the right. MyMPP: myeloid multipotent progenitor; LMPP: lymphoid-primed multipotent progenitors; LyMPP: lymphoid multipotent progenitor; LT-HSC: long-term hematopoietic stem cell; MEP: megakaryocyte/erythroid progenitor; CDP: common dendritic cell progenitor. (**F**) Cell density plots of the HSPC compartment in control or exercised young and old mice. (**G**) UMAP of subclusters in MuSCs. Fraction of each cluster is shown on the right. (**H**) Heatmap of marker genes for the MuSC clusters.

We annotated cell clusters based on the expression levels of canonical cell-type-specific markers in each tissue (Figures S1A-S1D). Notably, while *Pecam1*^+^ endothelial cells (ECs) and *Acta2^+^* smooth muscle cells from skeletal muscle and the SVZ (where they are referred to as mural cells) formed distinct clusters, B cells, T cells, monocytes and macrophages from all 4 tissues clustered together (Figure 1B). In addition to these immune cells, natural killer (NK) cells were also found in abundance in peripheral blood (Figure S1E). All annotated cell types were present in the 4 experimental groups, control young mice (Y-C), exercised young mice (Y-Ex), control old mice (O-C) and exercised old mice (O-Ex). Among all cell types, activated neural stem cells (aNSCs) and neuroblasts exhibited the most significant reduction with age, and exercise expanded these two populations of cells in both young and old animals (Figures 1C and 1D), supporting previous reports of the beneficial effect of exercise on adult neurogenesis (van Praag et al., 1999; van Praag *et al*., 2005). In the HSC compartment, aging was associated with an expansion in the long-term HSCs (LT-HSCs) (Figure 1E). Exercise was associated with a shift of cell expansion toward the myeloid multipotent progenitors (MyMPPs) in both young and old animals (Figure 1F). The vast majority of MuSCs in adult animals reside in quiescence in skeletal muscle. Although neither aging nor exercise significantly altered the cell cycle profiles or the relative ratio of MuSCs among all cell types in the muscle (Figures S1F and S1G), we found that subsets of MuSCs were selectively affected by aging or exercise (Figure 1G). These subsets of MuSCs were not defined by their differential expression of myogenic genes but by genes in metabolic and immune functions (Figures S1G and 1H). The subset of MuSCs that diminished in aging animals exhibited a gene expression signature of glycolysis (cluster 1) and mitochondrial respiration (cluster 3). The subset of MuSCs that were affected by exercise exhibited a gene expression signature that indicated a change in immune response (cluster 0 and 1). MuSCs in cluster 0 reduced in number in response to exercise regardless of the age of the animals, and those in cluster 1 reduced in number in O-C animals but was restored in O-Ex animals, suggesting that exercise may restore glycolysis in MuSCs by altering the immune profile of these cells. This finding provides a link between the regulation of metabolism in stem cells and the inflammatory environment in stem cell niche. Taken all three stem cell compartments together, these data indicate that aging leads to a change in the number or the relative ratio of subsets of stem cells, and exercise partially reverse cell composition changes in old mice.

### Effect of aging on the transcriptomes of stem cells and niches cells

To unveil common and unique gene expression changes associated with aging and exercise in skeletal muscle, brain, and the hematopoietic system, we identified differentially expressed genes (DEGs) from 22 major cell types in young control (Y-C), old control (O-C), young exercise (Y-Ex) and old exercise (O-Ex) mice (Table S1). Upregulated and downregulated DEGs in O-C when comparing to Y-C mice are defined as “age-DEGs”. We found that cells in the muscle were less impacted by aging based on the number of age-DEGs in each cell type (Figure 2A). Endothelial cells, smooth muscle cells (SMCs), monocytes, B and T cells from muscle all manifested smaller numbers of age-DEGs compared to their counterparts in other tissues. The most striking example was monocytes. We identified 960 age-DEGs in the monocytes from the blood, the most among all 22 cell types. In contrast, we identified 257 age-DEGs in the monocytes in aged muscle. Common age-DEGs in the same cell types between different tissues varied from 7.3% to 21.8% (Figure 2B). However, despite the distinct identity of age-DEGs in the same cell type from different tissue, these age-DEGs were implicated in similar biological pathways as exemplified by endothelial cells from the muscle and the SVZ (Figure S2A).

**Figure 2:**
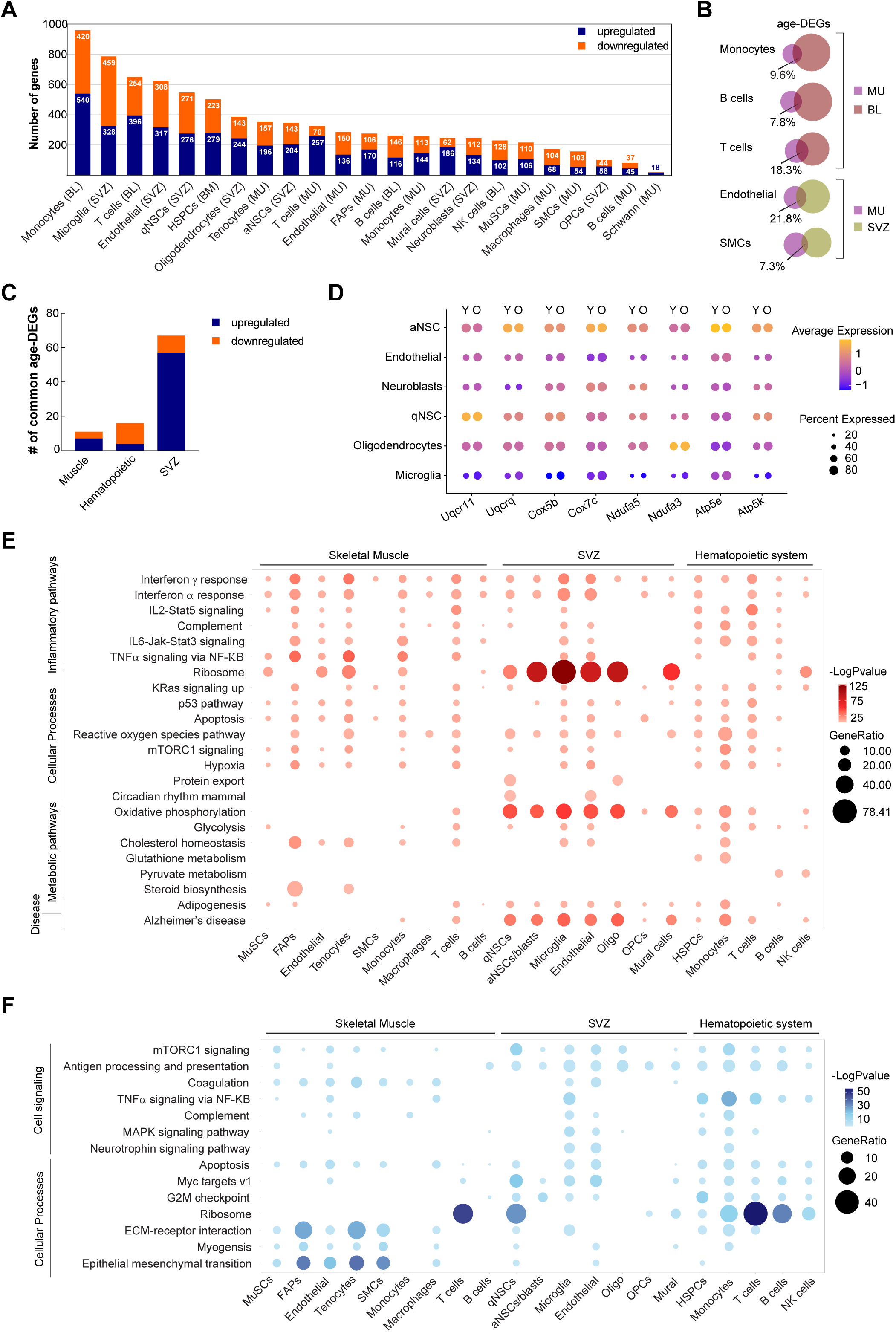
Effect of aging on the transcriptome of stem cells and niches cells. (**A**) Summary of the number of upregulated and downregulated age-DEGs in major cell clusters in all 3 tissues. Cell types are ranked by the total number of age-DEGs. **(B)** Venn diagrams demonstrating the percentage of common age-DEGs in specific cell types from different tissues. The size of the circles is proportional to the number of age-DEGs. **(C)** The number of common age-DEGs in the 3 stem cell compartments. **(D)** Dot plot demonstrating the expression of selected genes encoding enzymes involved in oxidative phosphorylation in SVZ cells from young and old animals. **(E)** Dot plot summarizing common biological pathways enriched among upregulated age-DEGs in major cell types of muscle, the SVZ, and the hematopoietic system. **(F)** Dot plot summarizing common biological pathways enriched among downregulated age-DEGs in major cell types of muscle, the SVZ, and the hematopoietic system.

To gain insight into the common and unique genes impacted by the aging process in these tissues, we separately compared upregulated and downregulated age-DEGs in HSCs and in major immune cell types in peripheral blood, muscle, and the SVZ (Fig. 2C). Common age-DEGs of cells in the muscle or the hematopoietic system did not show an enrichment of any pathways or biological processes. Among the three stem cell compartments, the major cell types in the SVZ had the highest number of common age-DEGs (Figures 2C and S2B). Among the 57 upregulated age-DEGs, 17 encode subunits of enzymes that play crucial roles in oxidative phosphorylation (Figures 2D and S2B). These enzymes, including ubiquinol-cytochrome c reductase (UQCR), cytochrome c oxidase (COX), NADH:ubiquinol oxireductase (NDUF) and ATP synthase, have been implicated in a number of neurodegenerative diseases such as Parkinson’s and Alzheimer’s disease (Manczak et al., 2004). In addition, 20 genes encoding ribosomal proteins or ribosome-associated proteins were found upregulated in SVZ cells from aged animals, suggesting a change in proteome homeostasis in these cells. It is worth noting that although no one single gene changes expression across all cell types in these three tissues, genes encoding various members of the heat shock protein 90 family were consistently found in the common downregulated age-DEGs (Figure S2C). Given the role of heat shock proteins as chaperons for protein folding, this finding suggests that deterioration in proteome homeostasis may be a general mechanism that contribute to cellular aging.

We have also explored the biological implication of age-DEGs from the major cell types in skeletal muscle, the SVZ, and peripheral blood/bone marrow using gene set enrichment analysis (GSEA). We found that although different cell types from different tissues exhibited a high degree of variability in age-DEGs, the cellular functions and responses in which these DEGs are implicated converge into a set of specific GSEA hallmark and KEGG pathways. The upregulated age-DEGs across cell types clearly revealed that aging is associated with an elevated inflammatory systemic environment. The age-DEGs from all cell types in our analyses indicated an increase in the interferon gamma (IFNγ) response, and the age-DEGs from a majority of the cell types indicated an increase in the interferon alpha (IFNα) response (Figure 2E). The majority of cell types from the muscle and peripheral blood/bone marrow exhibited an elevated response in interleukin 2 (IL2), IL6 and tumor necrosis factor α (TNFα) signaling which were absent in the majority of SVZ cells. Strikingly, the expression of ribosomal proteins exhibited noteworthy changes with age. Significant upregulation of ribosomal genes with age was found in the majority of SVZ cells and in the majority of non-immune cells in skeletal muscle, whereas immune cells as well as HSPCs downregulated the expression of ribosomal genes with age (Figure 2F). Metabolically, a significant increase in oxidative phosphorylation and, to a less extent, glycolysis with age was observed in SVZ cells but not muscle cells. Similarly, a significant decrease in MYC target gene expression and G2M checkpoint genes with age was found in the SVZ and hematopoietic system but not in muscle, including stem cells in these tissues, consistent with changes in the cell number of NSCs and HSPCs but not MuSCs. Changes unique to muscle resident cells also included a decrease in the interaction with extracellular matrix (ECM) (Fig. 2F).

### Reversal of age-induced gene expression changes by exercise

We next explored the effect of exercise on gene expression patterns of various cell types across tissues. Upregulated and downregulated DEGs in exercised mice, compared to control mice of the same age, are defined as “exercise-DEGs” (Ex-DEGs). A much smaller set of Ex-DEGs were identified in most of the cell types from old animals compared to young animals, with notable exceptions in immune cells from muscle (Figure 3A). Among the stem cell types, HSCs exhibited the smallest set of Ex-DEGs. A higher number of Ex-DEGs were induced in various immune cell types in the muscle, whereas a much lower number of Ex-DEGs were induced in the immune cells in peripheral blood from old animals. In young animals, monocytes from peripheral blood exhibited the largest set of Ex-DEGs. In old animals, monocytes from muscle exhibited the highest number of Ex-DEGs which included 77% of Ex-DEGs identified in muscle monocytes from young mice (Figure S3A). Common Ex-DEGs in the same cell types from different tissues varied from 2.6% to 8.5%. The low percentage of common Ex-DEGs of cells from different tissues suggest that cellular response to exercise is largely affected by local signaling in each tissue.

**Figure 3:**
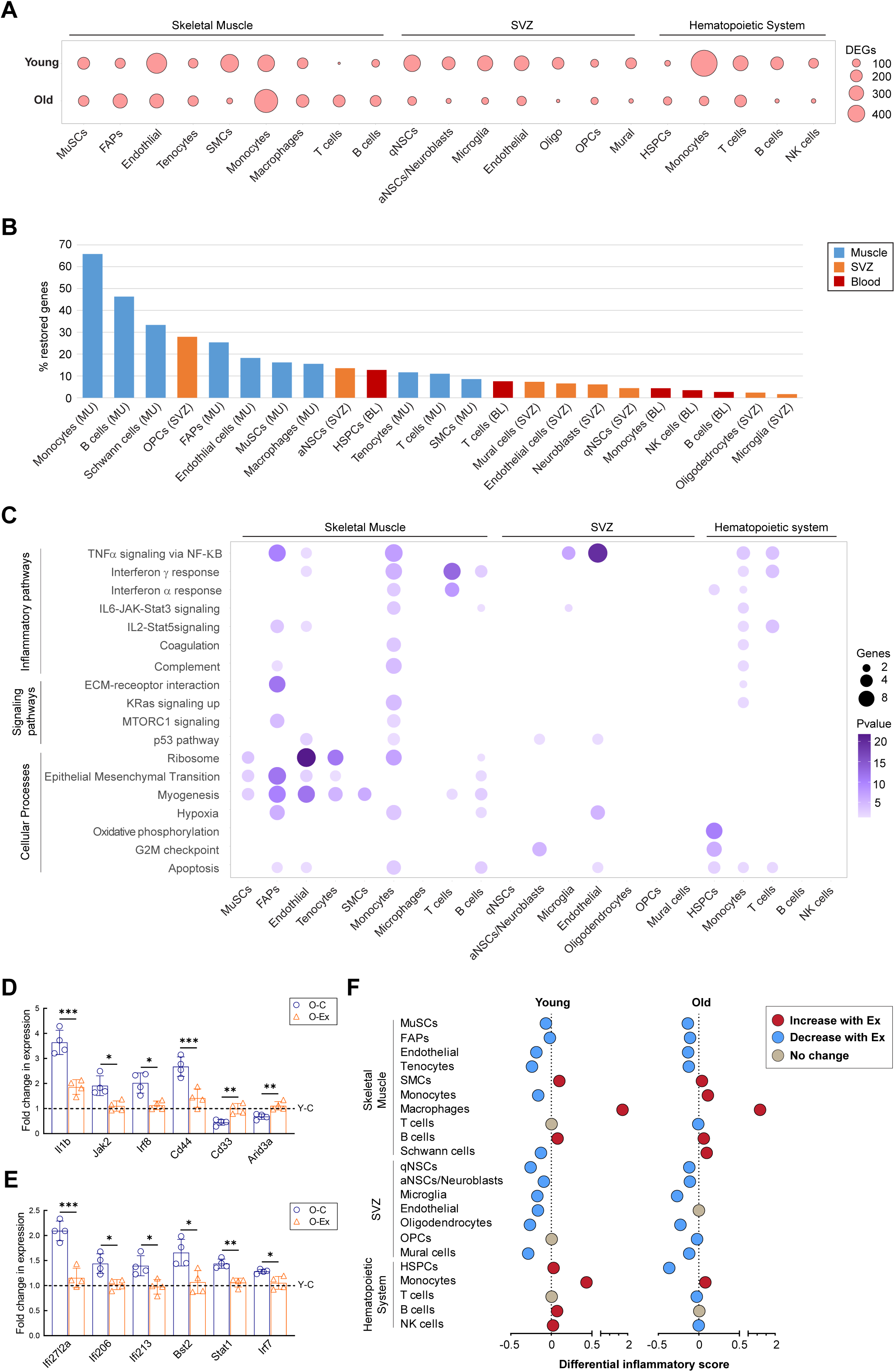
Reversal of age-induced gene expression changes by exercise. (**A**) Dot plot summarizing the number of Ex-DEGs in major cell types of muscle, the SVZ, and the hematopoietic system. **(B)** Bar graph demonstrating the percentage of exercise-restored genes in major cell types of the three tissues. Cell types are ranked from the highest percentage to the lowest. **(C)** Dot plot summarizing the biological pathways enriched among exercise-restored genes. **(D)** Bar graph demonstrating the relative expression levels of genes implicated in the TNFα pathway in muscle monocytes from old control and old exercised mice determined by RT-qPCR. For each gene, the fold changes in expression in comparison to the level in young control animals (Y-C, indicated by the dotted line) were plotted. **(E)** Bar graph demonstrating the relative expression level of genes implicated in the IFNγ pathway in muscle T cells from old control and old exercised mice determined by RT-qPCR. For each gene, the fold changes in expression in comparison to the level in young control animals (Y-C, indicated by the dotted line) were plotted. **(F)** Summary of differential inflammatory score in response to exercise in various cell types from young and old mice. Data are shown as mean ± SEM. *p < 0.05, **p<0.01, ***p < 0.001 (unpaired t tests).

We then performed pathway analyses with Ex-DEGs and found that exercise induced genes encoding metabolic enzymes implicated in oxidative phosphorylation in the majority of cells in young animals, including all cell types in the SVZ. In contrast, upregulation of these genes was almost absent in old animals (Table S1). Our analyses also suggested that a number of cytokine signaling pathways were suppressed by exercise (Table S2). In young animals, the downregulation of genes implicated in TNFα signaling was found in all non-immune cell types in muscle, all hematopoietic cells, and endothelial cells in the SVZ, most of which also exhibited evidence of downregulation in TNFα signaling in old animals. Genes implicated in IFNα and IFNγ signaling were also found to be downregulated in some cell types from young animals following exercise. Interestingly, suppression of IFNα and IFNγ signaling was found in immune cell types in muscle with exercise only in old animals.

We next investigated the effect of exercise on age-DEGs in these cell types. Downregulated age-DEGs that increase expression and upregulated age-DEGs that decrease expression in response to exercise are collectively termed “restored genes”. We found that cells in the muscle exhibited higher ratio of restored genes in comparison to cells from the other tissues (Figure 3B). Pathway analysis of restored genes revealed that nearly all cell types in muscle exhibited reversal of cellular signaling pathways and processes that changed during aging (Figure 3C). While the restored genes in non-immune cell types primarily related to tissue specific functions as exemplified by the enrichment of restored genes in pathways of epithelial-to-mesenchymal transition and myogenesis, restored genes identified in immune cells in muscle were enriched in cytokine signaling pathways. Consistent with the low percentage of restored genes in the majority cell types in the SVZ and hematopoietic systems (Figure 3B), exercise restored functional pathways that were altered in aged cells only in a limited number of SVZ and hematopoietic cell types (Figure 3C). Despite the fact that IFNα signaling and IFNγ signaling were found to be elevated in virtually all cell types with age (Figure 2E), exercise restored IFN signaling only in immune cell types in old animals (Figure 3C). On the contrary, TNFα signaling was restored by exercise in cell types in all three tissues which exhibited changes in this signaling pathway during aging. Using RT-PCR, we have confirmed the downregulation of target genes of the TNFα pathway in muscle monocytes (Figure 3D) and of the IFNγ pathway in muscle T cells (Figure 3E) in O-Ex in comparison to O-C animals.

To determine whether the effect of exercise on regulating the cellular inflammatory state was limited to the abovementioned pathways, we calculated an inflammatory score for each cell type based on the expression of selected genes in the GO term “inflammatory response”(Tirosh et al., 2016), and compared this score of each cell type between the experimental conditions. We found that in young mice, exercise was associated with a lower inflammatory score in nearly all cell types in the muscle and the SVZ, with the notable exception of macrophages from the muscle (Figures 3F and S3B-S3D). In old mice, the correlation between a lower inflammatory score and exercise was still present in most of these cell types albeit the difference was smaller than that in the young animals. In addition, although no difference was found in the inflammatory score of HSPCs in young animals, the inflammatory score of HSPCs in the O-Ex mice was lower than that in the O-C mice. These analyses suggest that exercise has a broad effect on the overall inflammatory landscape of stem cells and their niche environment.

### Effect of exercise on immune cells

In adult animals, monocytes and macrophages derive from MyMPPs, which are themselves progeny of LT-HSCs. The inclusion of all these cell types in our study allowed us to explore the link between the changes in the inflammatory score of HSPCs and the dichotomy between muscle monocytes/macrophages and monocytes from the peripheral blood in response to exercise in old animals. Towards this end, we first performed unsupervised clustering analysis on all monocytes/macrophages from muscle and blood in conjunction with LT-HSCs and MyMPPs. Using a resolution of 0.2 in Seurat, we identified 20 clusters in the monocyte/macrophage population (Figures 4A and 4B). We then performed trajectory analysis to infer the progression from LT-HSCs to various monocyte/macrophage clusters. By specifying the LT-HSCs as the root cell type, we established a trajectory along which clusters 3, 7, 9 and 15 were exclusively found in muscle (Figures 4C and S4A). Among these four clusters, 3, 7 and 9 were the most related based on their gene expression patterns (Figure S4B). Based on the trajectory, clusters 3 and 9 bifurcated from cluster 7 (Figure 4C). Cluster 3, situated at the end of a branch, likely represents a terminally differentiated monocyte/macrophage population in the muscle, whereas clusters 7 and 9 likely represent cycling monocytes given their transition state between the progenitors and other clusters of blood monocytes. We calculated the ratio of each cluster to the total pool of monocytes/macrophages, and found that despite an overall increase in monocyte number in both aged and exercised animals (Figure S4C), cluster 3 exhibited a significant reduction not only in cell number but also in relative fraction in all monocytes in O-C in comparison to Y-C mice, and the number of cells in this cluster appeared to increase in O-Ex mice (Figures S4C and S4D). This is further supported by the expression pattern of differentially expressed genes of these clusters in the UMAP space, where hematopoietic cell lineage genes (*Cd7, Flt3, Anpep, Itga3*) were found to be enriched in cluster 7, and functional phenotype-related genes, including *C1qa*, *C3ar1*, and *Plau*, which encode complement coagulation cascade proteins, including *Colec12* and *Itga5*, which encode phagosome proteins (Figure 4D).

**Figure 4:**
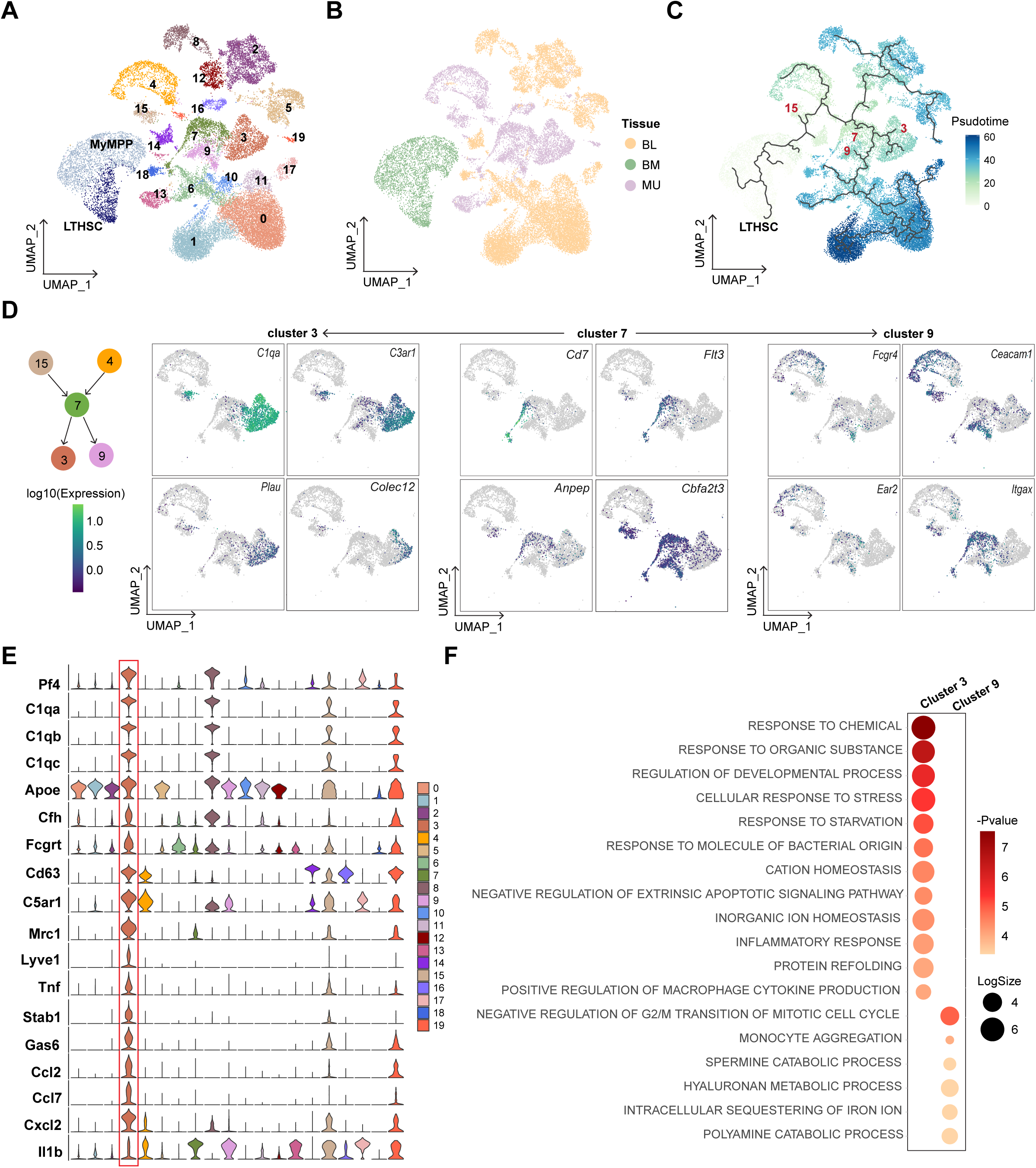
Aging and exercise-induced changes in immune cells. (**A-C**) UMAPs of HSCs, MyMPPs, and all monocytes/macrophages indicated by Louvain clusters, by tissues, and by Pseudotime trajectory, respectively, using HSCs as the root cell type, **(D)** UMAPs showing the expression pattern of specific genes in clusters 3, 7, and 9 monocytes/macrophage. **(E)** Violin plots showing the expression of monocyte/macrophage cluster 3 marker genes. **(F)** Dot plot summarizing the GO terms enriched among marker genes of clusters 3 and 9 of monocytes/macrophages.

We then identified marker genes of each monocyte/macrophage cluster (Table S3). To gain insight into the distinct features of cluster 3 that separated cells in this cluster from those in clusters 7 and 9, we compared the marker genes of these three clusters to signature gene sets of monocyte-derived macrophages of distinct phenotypes identified in a trajectory analysis of macrophages involved in skin wound healing (Sanin et al., 2022). We found that marker genes of cluster 3 included the highest expressing signature genes (*Pf4*, *C1qa*, *C1qb*, *C1qc*, *Apoe*, *Cfh*, and *Fcgrt*) as well as genes encoding surface markers (*Cd63* and *C5ar1*) of a class of phagocytic and remodeling macrophages (Figure 4E). Marker genes of cluster 3 also include *Mrc1*, which encodes CD206, often used as a marker of the tissue remodeling M2 macrophages, and a number of genes encoding cytokines known to have anti-inflammatory function. Marker genes of cluster 9 were found to be shared with other clusters of monocytes/macrophages, further supporting that these cells share a transition state with blood monocytes (Figure S4E). We performed FACS to enrich CD206^+^ macrophages and confirmed that they indeed express higher levels of *Pf4*, *Stab1*, *Gas6*, and *Cxcl2*, and lower levels of *Il1b* than CD206^-^ macrophages (Figures S4F and S4G). We have also performed gene ontology (GO) analysis on marker genes of clusters 3 and 9 to understand their functional differences. The analysis revealed that cluster 3 monocytes/macrophages expressed many genes involved in cellular responses to changes in various extracellular materials, including inorganic ions, organic nutrients, and microbial components (Figure 4F). In addition, these cells may play a role in the positive regulation of cytokine. Taken together, these analyses suggest that aging may be associated with the selective depletion of a subset of muscle resident monocytes that exhibit features of M2 macrophages, and exercise is associated with an increase in the number of these cells. The cytokines secreted by these macrophages may contribute to the changes in local stem cell milieu in response to exercise.

### Cell-cell communication network in skeletal muscle

To shed light on how the changes in muscle resident monocytes impact MuSCs and other niche cell types, we took advantage of the large number of cells and the sequencing depth in our data and generated cell-cell communication networks in muscle and the SVZ. To do this, we used the scRNA-seq analysis package CellChat which references DEGs to a comprehensively curated database of signaling molecules to identify overexpressed ligands and receptors for each cell group (Jin et al., 2021). A communication probability is then calculated based on the average expression of a ligand and its receptor by various cell groups. Significant interactions are identified based on random permutation of cell groups followed by recalculation between each pair of cell types. By applying this method, we identified a total of 65 potential signaling pathways mediated by secreted or cell surface signaling molecules, and 8 potential pathways mediated by extra-cellular matrix (ECM) components (Figures 5A and 5B) in skeletal muscle of young animals. A number of signaling pathways have been implicated in regulating MuSC homeostasis and function but the sources of ligands for these pathways thus far remain incompletely defined (Conboy et al., 2003; Pawlikowski et al., 2017; Ratajczak et al., 2003). Our analysis revealed the identity of cells that secret many of these ligands. For example, the ligands for the NOTCH pathway were contributed by endothelial cells, FGF signaling was from FAPs, Oncostatin M (OSM) signaling was initiated by monocytes, and CXCR signaling was initiated by SMC (Figure S5A). Interactions between each two-cell pairs in the muscle of young mice are summarized in Table S4.

**Figure 5:**
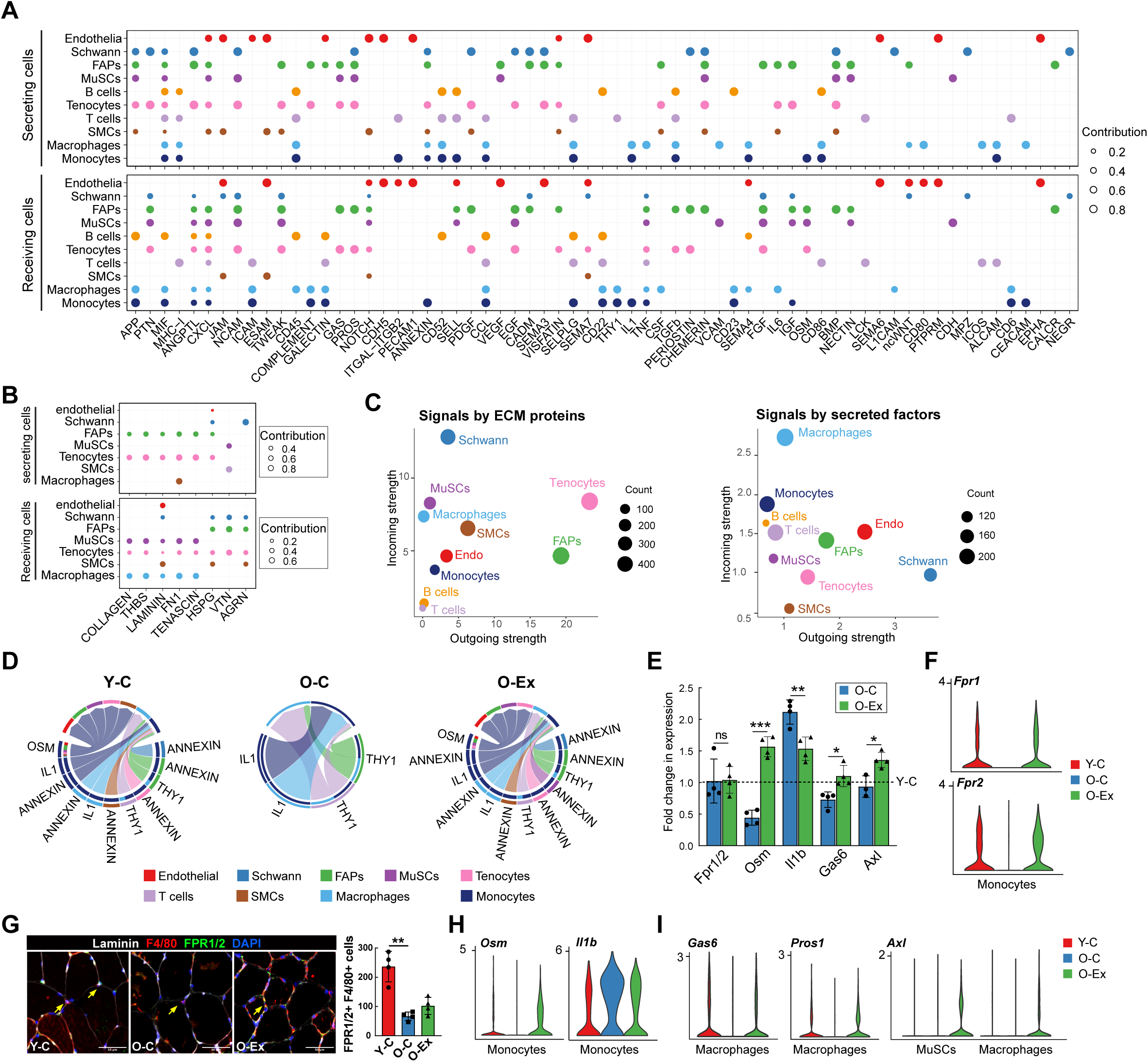
Cell-cell communication in the MuSC niche. (**A**) Dot plot summarizing the signal sending and receiving cells for each listed pathway mediated by secreted or cell surface molecules in skeletal muscle. The size of the dots is proportional to the contribution of a cell type to a specific pathway. **(B)** Dot plot summarizing the signal sending and receiving cells for each listed pathway mediated by ECM signaling in skeletal muscle. **(C)** Scatter plot showing the contribution of each cell type in the muscle to secreted/cell surface molecule-mediated communication (left) and to ECM-mediated communication (right). The size of the dots is proportional to the total number of incoming and outgoing signaling pathways associated with a cell type. **(D)** Chord plots showing cell-cell communication mediated by the ANNEXIN, OSM, IL1, and THY1 pathways in the muscle stem cell niche. The lower section of the circle represents signal sending cells and the top section of the circle represents signal receiving cells. At the lower section, the color bars at the outer circle represent signal sending cells, and those at the inner circle represent signal receiving cells. The arrows point to signal receiving cells. Note that ANNEXIN signaling and OSM signaling become undetectable with age but are restored by exercise. **(E)** Bar graph demonstrating the relative expression Fpr1, Osm, Il1b, Gas6, and Axl in muscle macrophages from O-C and O-Ex mice. For each gene, the fold changes in expression in comparison to the level in young control animals (Y-C, indicated by the dotted line) were plotted. **(F)** Violin plots showing the expression of *Fpr1* and *Fpr2* in muscle monocytes. **(G)** Representative images of muscle cross sections stained with Fpr1/2 antibodies. Monocytes/macrophages were co-stained with the F4/80 antibody and marked by the arrows. The number of total Fpr1/2 expressing monocytes/macrophages were quantified on the entire cross section and plotted in the bar graph shown on the right. **(H)** Violin plots showing the expression of *Il1b* and *Osm* in muscle monocytes. **(I)** Violin plots showing the expression pattern of *Axl*, *Gas6*, and *Pros1* in MuSCs and muscle macrophages. Data are shown as mean ± SD. *p < 0.05, **p<0.01, ***p < 0.001 (unpaired t tests).

In comparison to other cell types in muscle, FAPs and tenocytes express high levels of ECM proteins and contributed the most to interactions mediated by ECM components in the muscle (Figures 5B and 5C). Terminal Schwann cells, located at the neuromuscular junction, contributed the most to the intercellular interaction mediated by secreted or cell surface factors whereas macrophages and monocytes received the most secreted and cell surface signals (Figure 5C). With age, intercellular communication appeared to diminish in interactions initiated by Schwann cells, tenocytes, and T cells, and to a lesser degree, endothelial cells, whereas the interactions initiated by other cell types appeared generally enhanced (Figure S5B, left). Exercise restored the interaction initiated by Schwann cells, endothelial cells and FAPs (Figure S5B, right). In total, we identified 38 signaling pathways that changed with age, half of which were restored or partially restored after exercise (Figures S5C and S5D), including ANNEXIN, OSM, IL1 and GAS signaling that are primarily initiated by monocytes and macrophages (Figure 5D). Annexin 1 (ANXA1) is a phospholipid binding protein that has anti-inflammatory activity (Perretti and D’Acquisto, 2009). It is primarily produced by muscle monocytes/macrophages but also by other cell types (Figure S5C). Monocytes and macrophages express FPR1 and FPR2, which are receptors for ANXA1. We found no reduction in Fpr1 or Fpr2 expression in macrophages by RT-PCR (Fig. 5E). However, the number of Fpr1/2-expressing macrophages was significantly reduced in muscle cross sections from O-C mice, and the muscle of O-Ex mice exhibited a modest increase in the number of these cells (Figures 5F and 5G). The change in OSM and IL1 signaling was due to changes in both the number of macrophages that express the ligands and the levels of their expression in these cells (Figures 5E and 5H), whereas the change in GAS signaling was due to a change in the expression of the ligand Gas6 as well as the receptor Axl (Figure 5I). It is worth noting that exercise was associated with Axl expression also in MuSCs in old animals. As the receptor tyrosine kinase Axl has been recently shown to be associated with the self-renewal of human mammary epithelial progenitors (Engelsen et al., 2020), it will be interesting to determine whether MuSCs expressing Axl exhibit improved self-renewal potential in exercised old mice. Taken together, these findings suggest that exercise has an immune modulating effect in skeletal muscle, and intercellular interactions in the MuSC niche were largely restored to youthful levels by exercise.

### Cell-cell communication network in the SVZ

We have also performed cell-cell interaction analysis in the SVZ. In young animals, we identified a total of 52 potential signaling pathways mediated by secreted or cell surface signaling molecules and six potential signaling pathways mediated by ECM components (Figures 6A and 6B). Interactions between each two-cell pairs in the SVZ of young mice are summarized in Table S5. OPCs mediated the most interactions by secreted and cell surface proteins, whereas mural cells are the major initiator of ECM interactions (Figure 6C). Strong interactions and a high degree of signaling similarity were found between the cell types of ectodermal origin including qNSCs, aNSCs, neuroblasts, OPCs and oligodendrocytes, whereas cells of the mesodermal origin, including mural, microglia and endothelial cells, exhibit similar signaling patterns (Figure S6A). Although a high degree of signaling similarity was found between the neural cell types, we found that the qNSCs and aNSCs exhibited distinct signaling patterns with cells in the SVZ niche. For example, qNSCs but not aNSCs expressed WNT7b and interacted with the Frizzle receptors on aNSCs, OPCs and endothelial cells (Figure S6B), whereas aNSCs but not qNSCs expressed EFNA2 and EFNA5 and interacted with EPH receptors on themselves, OPCs, and, to a lesser degree, qNSCs, neuroblasts and endothelial cells. This cell-cell interaction analysis provides valuable insights into the common and distinct signaling pathways that regulate NSCs and various niche cell types.

**Figure 6:**
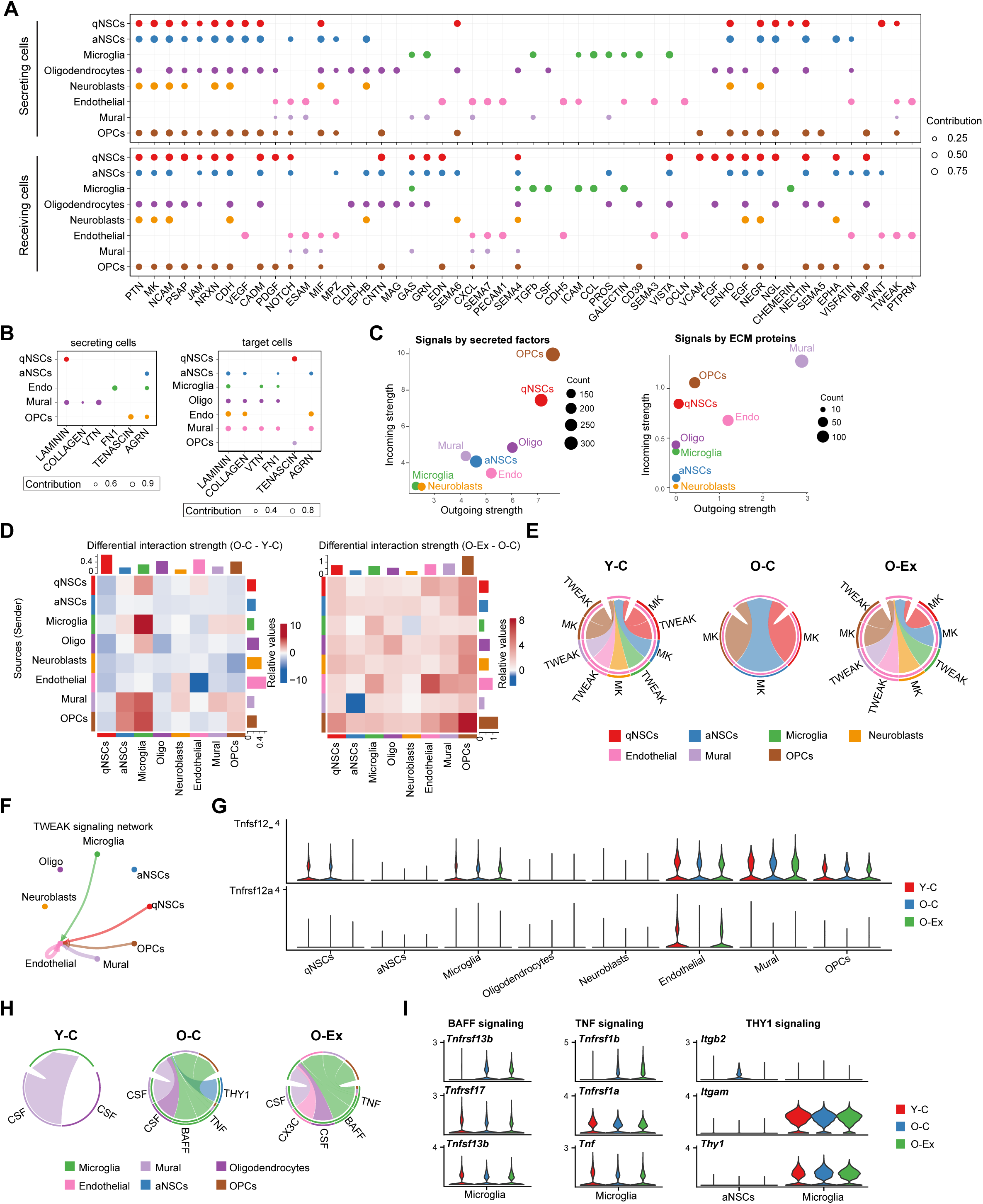
Cell-cell communication in the NSC niche. (**A**) Dot plot summarizing the expression pattern of secreted and cell surface signaling molecules and their receptors in the SVZ. The size of the dots is proportional to the contribution of a cell type to a specific pathway. **(B)** Dot plot summarizing the signal sending and receiving cells for each listed pathway mediated by ECM signaling in the SVZ. **(C)** Scatter plot showing the contribution of each cell type in the SVZ to secreted/cell surface molecule-mediated communication (left) and to ECM-mediated communication (right). The size of the dots is proportional to the total number of incoming and outgoing signaling pathways associated with a cell type. **(D)** Heatmaps showing the differential overall signaling strength in the SVZ between Y-C and O-C (left) and between O-C and Y-Ex (right) mice. The top bars and right bars represent the sum of incoming and outgoing signaling strength of each cell type, respectively. On the left, red and blue represent higher and lower signaling strength in the O-C mice, respectively, in comparison to Y-C mice. On the right, red and blue represent lower and higher signaling strength in the O-C mice, respectively, in comparison to O-Ex mice. **(E)** Chord plots showing the TWEAK and MK signaling network in the SVZ. Note that TWEAK signaling disappears in O-C and recovers in O-Ex. The arrows point to signal receiving cells. **(F)** Circle plot demonstrating the TWEAK signaling network in the SVZ. The thickness of the lines represents signaling strength; the color of the lines represents the source of the signal and the arrows point toward signal receiving cells. **(G)** Violin plots showing the expression pattern of the TNFSF12 ligand and the TNFRSF12A receptor in SVZ cells. **(H)** Chord plots showing the CSF, BAFF, TNF, THY1, and CX3C signaling networks in SVZ cells. **(I)** Violin plots showing the expression pattern of the BAFF, TNF, and THY1 signaling components in microglia.

Overall, aging is associated with a reduction in the intercellular interactions in the SVZ (Figure 6D, left). Neuroblasts and aNSCs exhibited a decrease in their interaction with all types of SVZ cells in old animals. This decrease is attributable to the depletion of aNSCs and neuroblasts with age (Figure 1D), as the cell communication calculation factors in the effect of population size on interaction strength. Consistent with this, the expansion of aNSCs and neuroblasts coincided with an increase in their interaction with all types of SVZ cells in O-Ex mice (Figure 6D, right). We identified 27 signaling pathways that changed with age in the SVZ, 13 of which were restored or partially restored after exercise (Figures S6C and S6D). Our analysis revealed that a number of cytokine signaling pathways change with age and exercise.

SVZ cell types interacted with endothelial cells via TWEAK signaling (Figures 6E and 6F). Aging was associated with a loss of *Tnfrsf12a* expression in endothelial cells and lack of TWEAK signaling in the SVZ (Figures 6E and 6G). Exercise was associated with the restoration of *Tnfrsf12a* expression in endothelial cells and thus TNFSF12 signaling (Figures 6F and 6G). TNFSF12 has been found to promote the proliferation and migration of endothelial cells and thus considered a pro-angiogenesis cytokine (Lynch et al., 1999; Wiley et al., 2001). Our findings suggest that it may be a key mediator of the beneficial effect of exercise on vascular health in the brain. On the contrary, the interaction between microglia and a number of SVZ cell types was enhanced in old nice (Figure 6D, left) and was further elevated by exercise (Figure 6D, right). Microglia in O-C animals gained active BAFF, THY1 and TNF signaling (Figure 6H). The activation of these pathways was due to the expression of genes encoding TNFSF13b and TNF by microglia and THY1 by aNSCs (Figure 6I). Exercise was associated with an abolishment of *Thy1* expression by aNSCs (Figure S6C). However, the expression of *Tnfsf13b* and *Tnf* in microglia was not affected by exercise in old animals. Interestingly, while TNFSF13b and TNF are generally believed to be pro-inflammatory, exercise of old animals was associated with the endothelial expression of the anti-inflammatory cytokine Cx3cl1 (Figure S6C), resulting in the activation of CX3C signaling in microglia. Active CX3CL1 signaling has been shown to protect neurons from IFN-induced cell death (Cho et al., 2011). Taken together, these data reveal potential mechanisms by which exercise mitigates the decline of brain health by reprogramming the cell-cell communication network in the neurogenesis region.

## Aging and Exercise-induced Changes in Myofibers

Not only are muscle fibers the structural basis for mobility during exercise, but they also secret circulating factors termed myokines that can affect distant tissues and cells (Pedersen and Febbraio, 2012). However, due to the multinucleated nature of muscle fibers, they are excluded from the scRNA-seq analysis. To understand the effect of aging and exercise on muscle fibers, we isolated individual muscle fibers free of attaching cells from the 4 groups of mice and performed RNA-seq analysis (Figure 7A). Although Y-C and O-C fibers were transcriptionally distinct, Y-Ex and O-Ex fibers shared a high degree of similarity in their transcriptomes (Figure 7A). DEGs of fibers are summarized in Table S6. The most prominent function of the DEGs that distinguished the fibers from exercised and control animals regardless of age was in the insulin signaling pathway (Figure S7A). Exercise led to changes in distinct metabolic and immune pathways in animals of different ages. In young animals, Ex-DEGs were implicated in the citrate cycle, unsaturated fatty acid biosynthesis, and IFNγ response, whereas in old animals, Ex-DEGs were implicated in glycerophospholipid metabolism and TGFβ signaling (Figure S7B). We found 24% of the downregulated age-DEGs restored in myofibers of O-Ex animals. The most prominent pathway that these restored genes implicated in was the lipid metabolic process (Figure 7B). Among the upregulated age-DEGs, 15% were restored, including those implicated in TNFα signaling pathway (Figure 7C).

**Figure 7:**
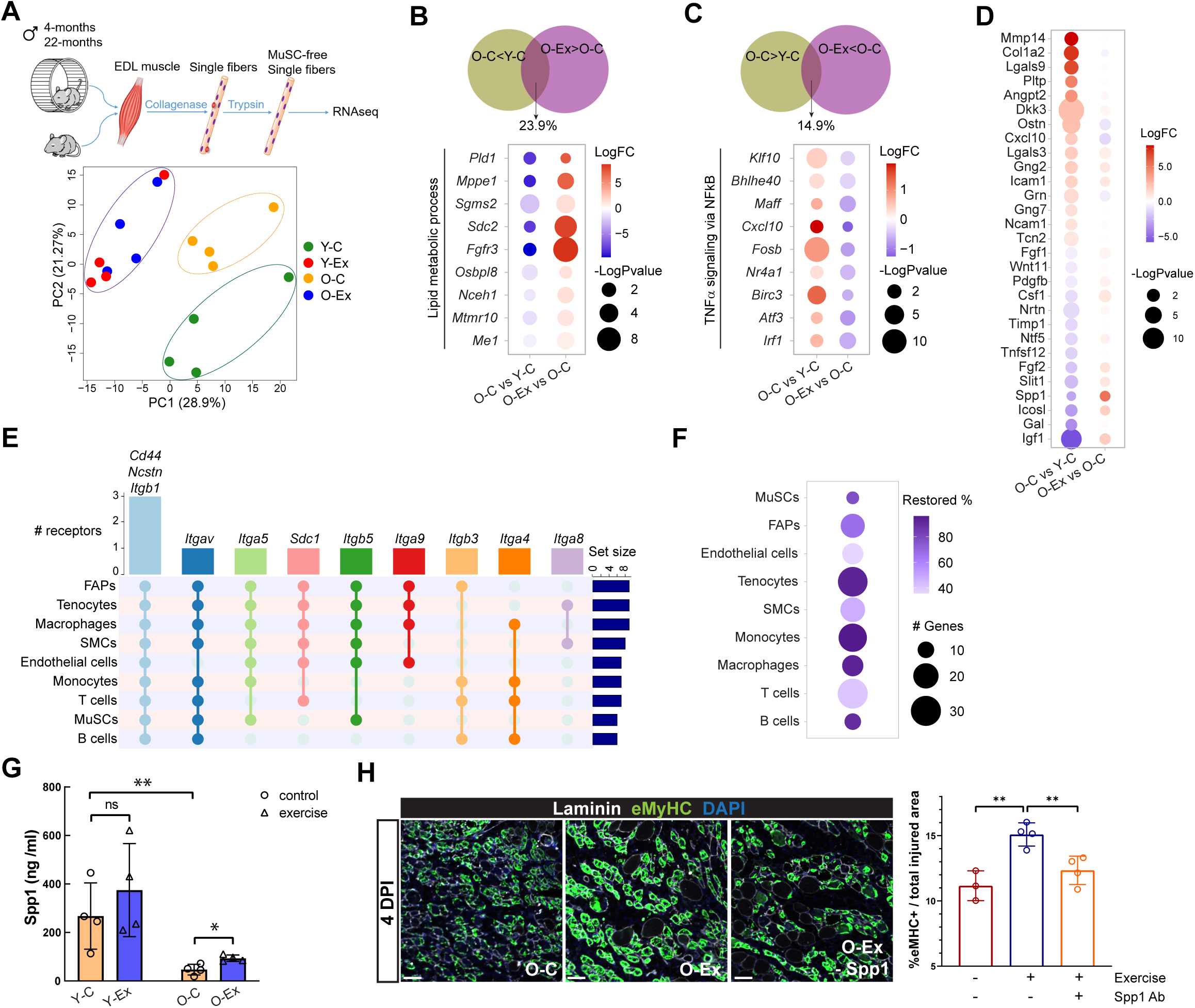
Aging- and exercise-induced changes in myofibers. (**A**) Schematic diagram of the single fiber RNA-seq methodology (top) and PCA analysis of data from fibers from young and old mice with and without exercise (bottom). **(B)** Venn diagram demonstrating the percentage of downregulated age-DEGs that were restored in myofibers from O-Ex mice (top) and dot plot summarizing the changes in the expression of genes in the lipid metabolic process (bottom). **(C)** Venn diagram demonstrating the percentage of upregulated age-DEGs that were restored in myofibers from O-Ex mice (top) and dot plot summarizing the changes in the expression of genes in the TNFα signaling pathway (bottom). **(D)** Dot plot demonstrating the differential expression of age-DEGs encoding secreted or cell surface ligands in myofibers from O-C mice in comparison to Y-C (left) and to O-Ex mice (right). The color indicates fold change and the size indicates the significance of the change. Red indicates higher and lower expression in O-C on the left and right, respectively. **(E)** UpSet plot showing the predicted receptors for Spp1 in cell types in skeletal muscle. **(F)** Dot plot demonstrating the number of predicted SPP1 target genes whose expression changed in old mice and was restored in exercised old mice, and the percentage of these genes among all SPP1 target genes whose expression changed in old mice. **(G)** Bar graph showing the level of Spp1 in Y-C, Y-Ex, O-C, and O-Ex mice detected by ELISA. (**H**) Comparison of muscle regeneration in O-C, O-Ex, and O-Ex mice injected with a neutralizing Spp1 antibody (O-Ex – Spp1). Cross sections were collected from the TA muscles of these mice 4 days after injury. Newly regenerated myofibers were identified by an antibody specific to embryonic myosin heavy chain (eMyHC). Scale bars represent 50 µm. The eMyHC-positive area normalized by total injured area is plotted and shown on the right. Data are shown as mean ± SEM. *p < 0.05, **p<0.01, ***p < 0.001 (unpaired t tests).

To determine how muscle fibers modulate the local niche environment for MuSCs and the systemic milieu that may signal distant tissues and cells, we compared the RNA-seq output data to a comprehensive database of ligand-receptor interaction (Shao et al., 2021), and we identified age-DEGs and Ex-DEGs encoding secreted proteins (Figures 7D and S7C). Exercise induced the expression of a larger number of secreted protein in young animals in comparison to old animals (Figure S7C). The fold change exhibited by these Ex-DEGs encoding secreted proteins was also greater in young animals. We found that exercise decreased the expression of cytokine genes that were upregulated with age such as *Cxcl10*, and increased the expression of a number of growth factors, such as Insulin Growth Factor 1 (IGF1), Secreted Phosphoprotein 1 (SPP1) and Fibroblast Growth Factor 2 (FGF2), which were downregulated with age (Figure 7D). The increase in *Spp1* expression in myofibers was consistent with our previous intercellular interaction analysis that SPP1 signaling was induced in nearly all types of muscle resident cells in O-Ex mice (Figure S5C). *Spp1* has been shown to be a downstream target of Peroxisome Proliferation Activator Receptor γ Coactivator 1α (PGC-1α) to activate macrophages and enhance angiogenesis in skeletal muscle (Rowe et al., 2014). We used the NicheNet cell-cell interaction package to identify receptors and target genes of SPP1 in muscle resident cells (Browaeys et al., 2020). Our analysis predicted that receptors of SPP1 are expressed in all major cell types in muscle (Figure 7E), and that exercise was associated with the reversal of the change in the expression of the majority of predicted SPP1 target genes across muscle cell types (Figures 7F and S7D, Table S7). These reversed SPP1 target genes include cytokine genes *Ccl11*, *Ccl3*, *Il1b*, *Csf1*, *Btg2*, and genes encoding transcription factors, such as *Irf8* and *Stat1*, which play important roles in regulating immune function (Figure S7D). We performed ELISA analysis to confirm the reduction of Spp1 levels in both skeletal muscle and plasma in aged animals and that exercise was associated with elevated Spp1 levels (Figures 7G and S7E). To determine whether Spp1 mediates the beneficial effect of exercise on muscle regeneration, we injured the tibialis anterior (TA) muscles of control or exercised old mice and harvested the injured TA muscles 4.5 days after injury to assess regeneration. We compared the area with regenerating muscle fibers as indicated by embryonic myosin heavy chain (eMyHC) expression from these mice. We found that exercise improved muscle regeneration in old mice, and the inhibition of Spp1 by a neutralizing antibody attenuated the effect of exercise (Figure 7H). Taken together, these data suggest that in response to exercise, myofibers produce SPP1 which may improve skeletal muscle homeostasis and regeneration by immune modulation in old animals.

## DISCUSSION

Tissue atlases at the single cell transcriptome level have been generated across the lifespan of mammalian species in recent years (Ma *et al*., 2020; Martinez-Jimenez *et al*., 2017; Tabula Muris, 2020). These atlases have provided unprecedented resolution on cell composition and transcriptomic changes in a broad range of tissues in aged animals. However, due to their rarity, stem cells are often not identifiable in these atlases, or are present at a frequency that is too low to reveal statistically significant changes between age groups. Our study set out to understand the cell type composition changes in stem cell compartments and the molecular changes that occur in stem cells and niche cells during aging and how exercise as an aging intervention modulates these changes. Towards this end, we generated a single cell transcriptomic atlas with sufficient representation of HSCs, MuSCs and NSCs, as well as their surrounding niche cells and/or progeny, from young and old mice subjected to voluntary exercise or conventional housing. This work complements existing atlases in expanding our knowledge on the systemic changes that occur during aging, and provides a blueprint to compare changes in the MuSC niches in muscles of different developmental origin and in the NSC niche in hippocampus in response to exercise. Exercise clearly elicits metabolic and endocrine changes. Although our study and analyses focuses on the effect of exercise on age-related changes in inflammation, changes in metabolic genes and pathways are also readily extractable from our atlas. As such, our work has the potential to be the foundation of many future studies to understand the mechanisms that mediate the effect of exercise on the functions of stem cells and their niche.

To ensure that the multinucleated myofibers, the cells responsible for locomotion, were included in our transcriptomic analysis, we also performed RNA-seq with single myofibers that are devoid of neighboring mononucleated cells. Accompanying the development of scRNA-seq technologies, computational tools have also been developed to infer changes in cell type composition and in the transcriptome of a defined cell type from whole tissue bulk RNA-seq data (Newman et al., 2019). The combination of our scRNA-seq with skeletal muscle, the single fiber RNA-seq and existing bulk RNA-seq data from the entire muscle provides an ideal dataset to develop and test such computational tools. Skeletal muscle is the largest organ of the mammalian body. In addition to the contractile function, increasing evidence has indicated that skeletal muscle is also a secretory organ in response to exercise (Pedersen and Febbraio, 2012). However, the exact cellular source for many exercise-induced cytokines and growth factors in muscle tissue remains unclear. The datasets generated in our study provide a foundation to distinguish myokines, defined as cytokines secreted by myofibers (Pedersen and Febbraio, 2012), from those secreted by muscle resident cells. Understanding the regulation of these secreted proteins will provide valuable insights in the molecular mechanisms by which myofibers mediate the effect of exercise on muscle resident cells and distal tissues.

Previous scRNA-seq studies have provided compelling evidence that mammalian aging is associated with alterations in the immune system and in inflammatory responses. Manifestations of such changes include increased transcriptional variability during naïve T cell activation, infiltration of immune cells into the tissues where they are absent in young animals, and clonal expansion of lymphocytes (Dulken et al., 2019; Martinez-Jimenez *et al*., 2017; Tabula Muris, 2020). Consistent with findings from these studies, our scRNA-seq data reveal an increase in the expression of IFN responsive genes in all cell types and an increase in the expression of TNFα target genes in the majority of cell types from aged animals. In exercised old animals, the increased expression of genes in the IFN pathway is reversed in many cell types in the muscle and blood, and the increased expression of genes in the TNFα pathway is reversed in various cell types of the muscle, SVZ and blood. Unlike cells from the muscle and blood, most cells in the SVZ of old animals, with the exception of microglia and endothelial cells, exhibit elevated IFN but not IL6 or TNFα signaling. We also find that the age-induced upregulation of TNFα and IL6 target genes in microglia can be rescued by exercise while the upregulation of IFN signaling persists. These analyses support the previous finding that infiltrating T cells in the brain secrete IFNγ (Dulken *et al*., 2019), and suggest that exercise does not lead to the elimination of these infiltrating T cells. Interestingly, while the induction of IFN signaling in microglia in aged animals is not ameliorated by exercise, the cell-cell network analysis of the SVZ reveals that the expression of the cytokine gene *Cx3cl1* is induced by exercise in old animals. Active CX3CL1 signaling has been shown to suppress neuronal death in co-cultures of neurons with IFNγ-stimulated microglia and in mouse models of Alzheimer’s disease (AD) (Bhaskar et al., 2010; Cho *et al*., 2011; Mizuno et al., 2003). Deficiency of Cx3cl1 expression has also been found in brain specimen from AD patients (Cho *et al*., 2011). Whether active CX3CL1 signaling mediates the beneficial effect of exercise in reducing the risk of neurodegenerative conditions, including AD, by limiting the neurotoxicity resulting from abnormal microglia activation requires further study. Nontheless, our study and findings provide strong evidence that exercise ameliorates the age-related increase of inflammation in multiple tissues.

Our analysis has revealed an important role of monocytes/macrophages as well as microglia in the SVZ in the inflammatory changes in aging and after exercise. It has been implicated recently that the mesenchymal cells in the muscle and adipose tissues are involved in the remodeling of extracellular matrix in response to exercise and high-fat diet(Yang et al., 2022). Our cell-cell communication network analysis has revealed that FAPs, the mesenchymal progenitor cells in skeletal muscle, are one of the most prominent cell types that mediate intercellular signaling via extracellular matrix. Studies of the signaling pathways between FAPs and monocytes/macrophages predicted in our analysis will provide insights on specific mechanisms by which FAPs affect the inflammatory landscape in old animals and in response to exercise.

In summary, our single-cell atlas, in combination with the single myofiber transcriptomic profiles, provides a rich source to study the changes in the cell type composition, transcriptional network, and communication patterns during aging and with physical exercise. Elucidating the mechanisms by which exercise modulates these intercellular and intracellular processes has great potential for the discovery of molecules and pathways that can be targeted therapeutically to reduce age-related diseases and to increase organismal health span.

## LIMITATIONS

The skeletal muscle of male and female mice are different in strength and fiber type composition which are affected by sex hormones. The change in the endocrine system of male and female mice during aging is likely to affect how skeletal muscle and the hematopoietic system respond to exercise. In addition, the duration, intensity, and type (such as endurance versus resistance training) of exercise may also have different impacts on stem cells and their niche, in particular in the muscle compartment. Due to the extraordinary cost of single cell RNA-seq studies, this study is limited to the use of male mice with a fixed aerobic exercise regime to minimize individual variability and to increase the likelihood of obtaining statistically significant findings. These data will provide a framework for future studies of comparing male and female animals for each of the stem cell compartments across age and in response to various type of exercises.

Due to the rarity of HSCs in bone marrow, we used antibodies to enrich for hematopoietic lineages prior to single cell capture. As such, the HSC compartment in our study does not contain non-hematopoietic lineage such as stroma cells. Ependymal cells, which have been reported to be numerous in the SVZ (Shook et al., 2012), were present in extremely low numbers in our atlas. It is possible ependymal cells are not efficiently loaded in droplets or are sheared in the Chromium cell capture device due to their large size. In the MuSC compartment, although we performed single fiber RNA-seq to compensate for the lack of these large and multinucleated cells in our scRNAseq atlas, the incorporation of RNA-seq data into the scRNAseq atlas requires additional computational processing.

## AUTHOR CONTRIBUTIONS

L.L. and T.A.R. conceived of the study and designed the experiments. L.L., M.T.B., J.M.R., S.K., A.L., M.M., C.M-R, J.K., H.I and M.J. carried out the experiments. L.L., M.T.B., J.M.R., S.K., L.T. and M.W. performed data analysis. J.C.W., M.A.G., A. B. and T.A.R. provided supervision and guidance. All authors participated in data interpretation. L.L. and T.A.R. wrote the manuscript with the input of all of the coauthors.

## ACKNOWLEDGEMENTS

This work used the Genome Sequencing Service Center by Stanford Center for Genomics and Personalized Medicine Sequencing Center, supported by the grant award NIH S10OD025212, and NIH/NIDDK P30DK116074. We thank all members of the Rando, Brunet and Goodell laboratories for helpful discussions. We thank Brandon Carter for assistance at FACS core at VAPAHCS, and the Stanford Genome Technology Center for assistance in high-throughput sequencing. J.MR. is a Gilliam Fellow of the Howard Hughes Medical Institute. This work was supported by grants from the NIH (R01DK092883 and P30CA125123) to M.A.G, by a Chan Zuckerberg Initiative award, a Simons Foundation grant, and a generous gift from Michele and Timothy Barakett to A.B., by support of the Glenn Foundation for Medical Research and grants from the Department of Veterans Affairs (BLR&D Merit Review) and the NIH (R37 AG023806, R01 AR073248, and TR01 AG047820) to T.A.R, and by the NIH grant P01 AG036695 to T.A.R., A.B. and M.A.G.

## SUPPLEMENTAL INFORMATION

## Materials and Methods

### Animals and treatments

Male C57BL/6J mice of 4- and 22-months of age were acquired from NIA and housed in the Veterinary Medical Unit at the Veterans Affairs Palo Alto Health Care System. Animal protocols were approved by the Institutional Animal Care and Use Committee. For voluntary wheel running, mice were housed individually for 5 weeks in polycarbonate cages with 12.7-cm-diameter wheels equipped with optical rotation sensors (Lafayette Instrument, 80820). Non-exercised control mice were housed in identical cages with the wheels removed.

### Cell isolation

Cardiac puncture was performed to collect blood from anesthetized mice using syringes and tubes treated to prevent coagulation. The blood was diluted with equal volume of Dulbecco’s phosphate-buffered saline (PBS) and overlaid onto Lymphoprep (Stem Cell Technologies). Plasma and white blood cells were collected after gradient centrifugation. Live cells were sorted in the presence of DAPI for single cell capture. SVZ neurogenic niches were collected and processed as previously described (Dulken *et al*., 2019). Briefly, after blood collection, mice were perfused with 15 ml of PBS with heparin sodium salt (50 U/ml) (Sigma-Aldrich, H3149-50KU) to remove residue blood. Microdissection of the SVZ was performed immediately following perfusion. The dissociated cells from the SVZ were then centrifuged through 22% Percoll (Sigma-Aldrich, GE17-0891-01) in PBS to remove myelin debris. After centrifugation, cells were filtered through a 35-μm snapcap filter (Corning, 352235), washed once with 1.5 ml of FACS buffer (HBSS (Thermo-Fisher, 14175103), 1% bovine serum albumin (Sigma, A7979), 0.1% glucose (Sigma-Aldrich, G7021) and spun down for 5 min at 300*g*. Cells were then resuspended in FACS buffer with 1 μg/ml propidium iodide (PI) (BioLegend, 421301). FACS sorting was performed on a BD FACS Aria II sorter using a 100-µm nozzle. Isolation of cells from skeletal muscle by sequential enzymatic and physical dissociation was performed as previously described (Liu et al., 2015). Briefly, hindlimb muscles were incubated with collagenase II and dispase sequentially before physical trituration. The resulting cell suspension was filtered through 40 µm cell sieges to remove myofiber debris. Cells were then sorted in the presence of DAPI using a 70 µm nozzle on a BD FACS Aria II sorter and live cells were collected for single cell capture. HSCs were enriched from the bone marrow as previous described (Challen et al., 2009). Briefly, bone marrow was isolated from femurs, tibiae and iliac crests. Side population staining was performed with Hoechst 33342 (Sigma). Magnetic enrichment was carried out on an AutoMACS using microbeads conjugated to antibodies recognizing biotin or mouse CD117 (all from Miltenyi Biotec). HSCs (CD45.2^+^, side population^KLS^, CD150^+^) were sorted in the presence of 1 μg/ml PI with a 70 µm nozzle on a BD FACS Aria 3.3 sorter.

### Single cell RNA sequencing

Single cells were captured in droplet emulsions using a 10x Chromium Controller (10x Genomics) with a target output of 10,000 cells per sample. scRNA-seq libraries were constructed as per 10x Genomics protocol using GenCode Single-Cell 3’ Gel Bead and Library V3 kit according to manufacturer’s instructions. The libraries were then pooled and sequenced on a NovaSeq6000 Sequencing System (Illumina) according to manufacturer’s instructions.

### scRNA-seq data processing

Sequences were demultiplexed using CellRanger version 2.0 (10x Genomics) with default parameters. Reads were aligned to the mm10plus genome using STAR. Gene counts were produced using HTSEQ with default parameters except “stranded” was set to false and “mode” was set to intersection-nonempty. Only cells with >500 genes, >1000 UMIs, and <10% mitochondrial reads were included in our analysis. Scaling, normalization, variable gene selection, dimensionality reduction, and clustering were performed with default settings using the Seurat package version 3.1 (Stuart et al., 2019). Cell types were assigned to each cluster using known marker genes. Clusters of doublets were identified and removed by careful manual inspection. DEGs were identified using the Seurat “FindAllMarkers” function. Cell density plots were generated by first using the “scanpy.t.embedding” function in the Scanpy Python toolkit to calculate the density of UMAP representation of the cell type of interest followed by using the “pl.embedding.density” function to plot the results. To calculate the inflammatory score of each cell type, we select genes in the GO term “inflammatory response” that are detected in our scRNAseq analysis and added a module in Seurat using the AddModuleScore function(Tirosh *et al*., 2016). The Differential Inflammatory Score was calculated by the subtraction of the inflammatory score of exercise mice from that of control mice of the same age. A negative value indicates a decrease and a positive value indicates an increase in the score in exercised animals. Intercellular interaction analysis was performed using the CellPhoneDB (Efremova et al., 2020) and CellChat (Jin *et al*., 2021) packages with standard parameters. Receptors and target genes for selected pathways were predicted using the NicheNet package (Browaeys *et al*., 2020).

### Single fiber isolation

Single muscle fibers were prepared essentially as described (Rosenblatt et al., 1995). Briefly, the *extensor digitorum longus* (EDL) muscles were carefully dissected from mice following euthanasia and digested in 2 mg/ml collagenase II prepared in Ham’s F10 with 10% horse serum at 37°C for 80 min with gentle agitation. The digested EDL muscles were then triturated with a wide bore glass pipet in 20 ml 10% horse serum in Ham’s F10 in a 10 cm tissue culture dish.

Individual fibers were washed 3 times in medium and incubated with 0.25% trypsin for 15 minutes. Trypsinized fibers were then rinse 3 times and centrifuged at 500*g* for 5 minutes. Fibers were then snap-frozen in liquid nitrogen and stored at -80°C until RNA extraction.

### RNA-seq

RNA was extracted from fibers using the RNeasy Micro Plus Kit (Qiagen) according to manufacturer’s instructions. Reverse transcription was performed with 10 ng RNA using the SMART-Seq v4 Ultra Low Input RNA Kit (Takara) according to manufacturer’s instructions. The cDNA was then sheared with a Covaris ultrasonicator and library constructions were performed with the Ovation Ultralow Multiplex system (NuGEN). Libraries underwent paired-end 101-bp sequencing on a HiSeq 4000 sequencer (Illumina) at the Stanford Genome Sequencing Service Center to a depth of 20-40 million reads. Reads were adapter- and quality-trimmed with trim_galore (http://www.bioinformatics.babraham.ac.uk/projects/trim_galore) prior to mapping to the mouse genome assembly with STAR (Dobin et al., 2013). The number of reads in each gene was summarized using the package FeatureCounts (Liao et al., 2014) and DEGs were detected using the DESeq2 R package (Love et al., 2014). Gene Set Enrichment Analysis (Subramanian et al., 2005) was performed using the MSigDB Hallmark gene set and KEGG gene set.

### RT-qPCR

Total RNA was extracted from cells using the RNeasy Micro Plus Kit (Qiagen) according to manufacturer’s instructions. Reverse transcription was performed with the High Capacity cDNA Reverse Transcription Kit (Applied Biosystems). Quantitative PCR was performed on LightCycler 480 (Roche) with matching SYBR Green master mix. Each measurement was performed in triplicate. Relative quantification of gene expression normalized to *Gapdh* was carried-out using the comparative C_T_ method.

### ELISA

To prepare muscle extract for ELISA analysis, gastrocnemius muscles were homogenized in RIPA buffer supplemented with protease inhibitors. The homogenates were centrifuged at 12,000 rpm, 4°C for 15 minutes. The supernatant was collected for analysis. Mouse Spp1(OPN) ELISA (Invitrogen) assays were performed according to the manufacturer’s instructions. Briefly, muscle extracts were diluted 100-fold, and plasma samples were diluted 250-fold using the dilution buffer supplied by the manufacturer. The assay plate was incubated sequentially with the diluted samples or standards for 2.5 hours, 100 µl biotin conjugate for 1 hour, and 100 µl of streptavidin-HRP solution for 45 minutes. All incubation was at room temperature and with gentle shaking. Each well was washed four times with wash buffer after each incubation. After the last wash, 100 µl of TMB substrate was added to each well. When the wells with most concentrated standard turn deep blue, 50 µl of stop solution was added to each well and the absorbance at 450nm was read immediately. The Spp1 concentration in assay samples was calculated according to the standard curve. The Spp1 concentration in gastrocnemius muscles was normalized to their weight.

### Analysis of muscle regeneration

Male C57BL/6J mice of 22-months of age were subjected to voluntary wheel running for 3 weeks before muscle injury was induced. To test the role of Spp1 in muscle regeneration, 50 µl 1.2% barium chloride (w/v in H_2_O) with or without 2 µg of Osteopontin neutralizing antibody (AF808, R&D Systems) was injected into a TA muscle. Injured TA muscles were collected 4.5 days after injury, embedded in OCT, and frozen in liquid nitrogen cooled isopentane. To assess regenerating myofibers, 10 µm cross sections were collected from the frozen TA muscles and stained with laminin (anti-rabbit Sigma L9393) and embryonic Myosin Heavy Chain (eMHC, DSHB F1.652) antibodies. Sections were imaged with a Nikon ECLIPSE Ti2 fluorescent microscope. Quantification of eMHC staining was performed in ImageJ. Percent eMHC area was calculated as sum of eMHC area per TA / (total TA area – uninjured area).

### Statistical analysis

All statistical analyses were performed using GraphPad Prism 8 (GraphPad Software). Significance was calculated using two-tailed, unpaired Student’s *t*-tests unless stated otherwise. Differences were considered to be statistically significant at the *P* < 0.05 level (*p < 0.05, **p<0.01, ***p < 0.001, ns: not significant).

## Supplemental Figure Legends

**Figure S1:**
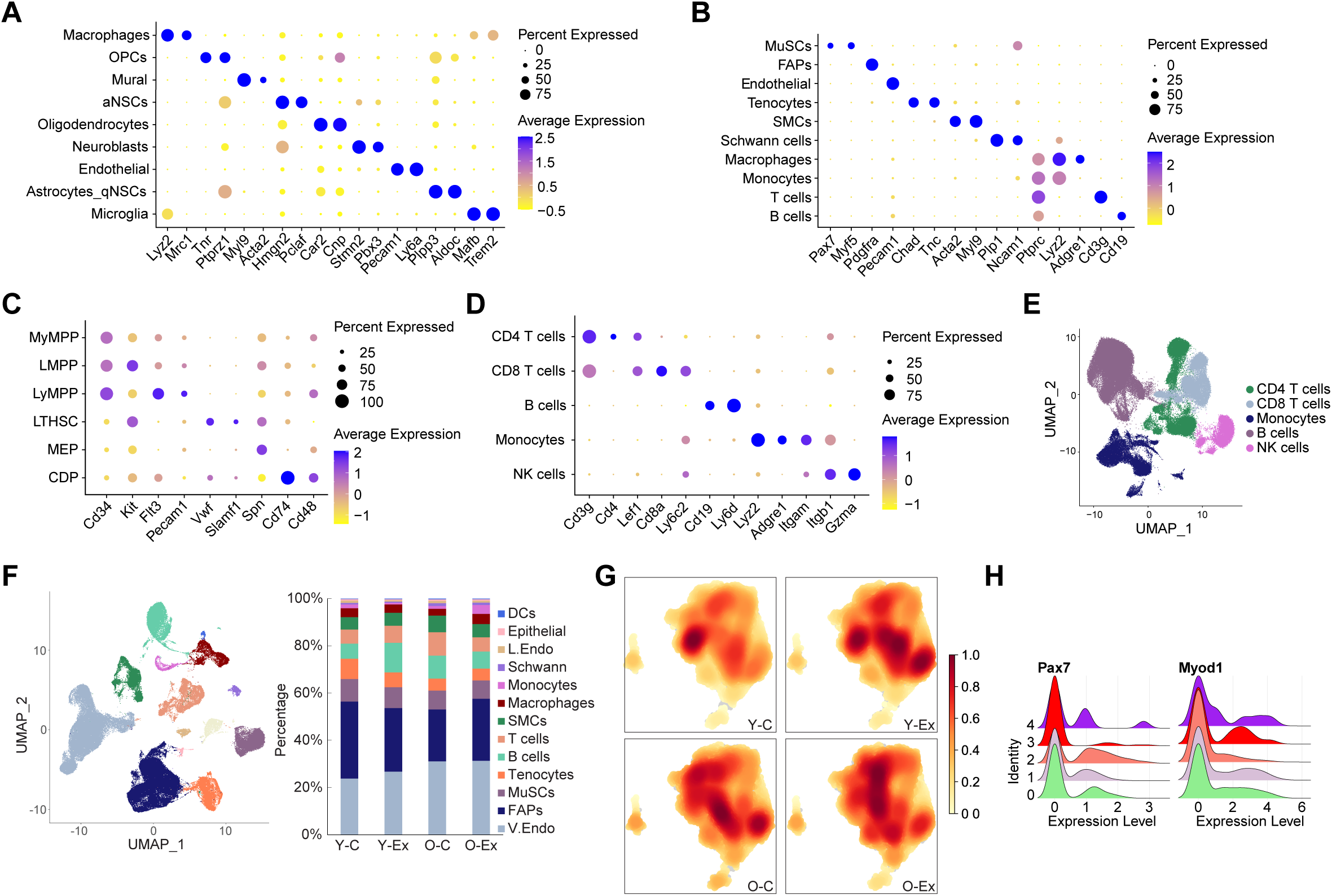
Annotation of cell types in aging and exercise mouse single-cell atlases. (**A-D**) Dot plot showing markers used for cell type annotation in the SVZ, in skeletal muscle, in the HSC compartment, and in blood immune cells. **(E)** UMAP showing major immune cell types in the blood. **(F)** UMAP of cell types in skeletal muscle. Fraction of each cell type is shown on the right. **(G)** Heatmap of cell density of MuSCs. (**H**) Ridge plots of the expression of myogenic markers Pax7 and Myod1 in the MuSC clusters illustrated in Fig. 1G.

**Figure S2:**
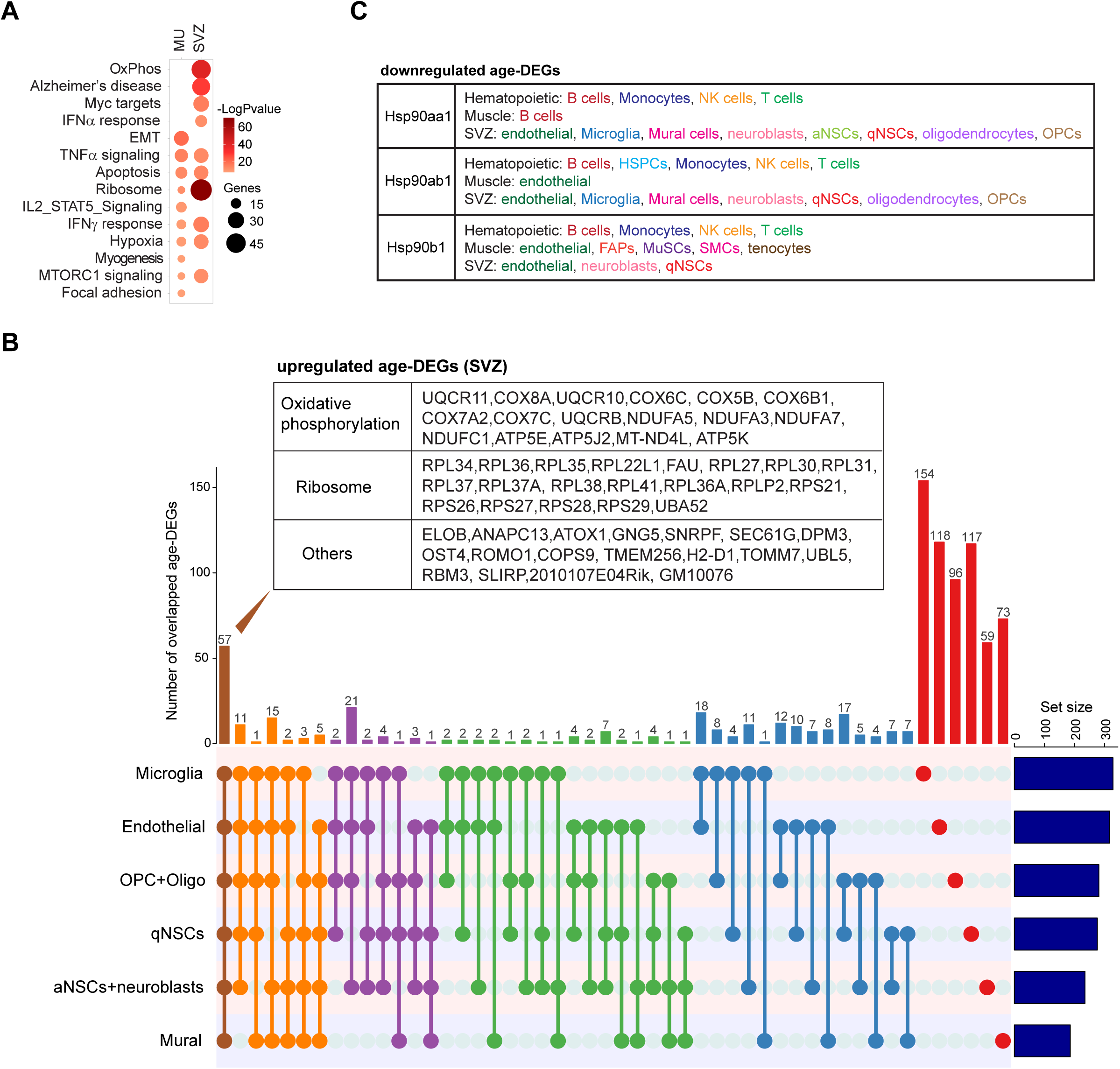
Common patterns of transcriptomic changes in stem cell compartments during aging. (**A**) Dot plot summarizing the top 10 (ranked by P value) biological pathways enriched among age-DEGs in endothelial cells of the muscle (MU) and the SVZ. **(B)** UpSet plot demonstrating common upregulated age-DEGs among major cell types in the SVZ. The age-DEGs are organized by the number of cell types in which they are found in common, with those expressed in all six cell types on the far left and those found in only one cell type on the far right. The common age-DEGs that were upregulated in all 6 types of cells are listed in the table. Oligo: oligodendrocytes. **(C)** Table listing cell types in which members of the Hsp90 family were downregulated with age.

**Figure S3:**
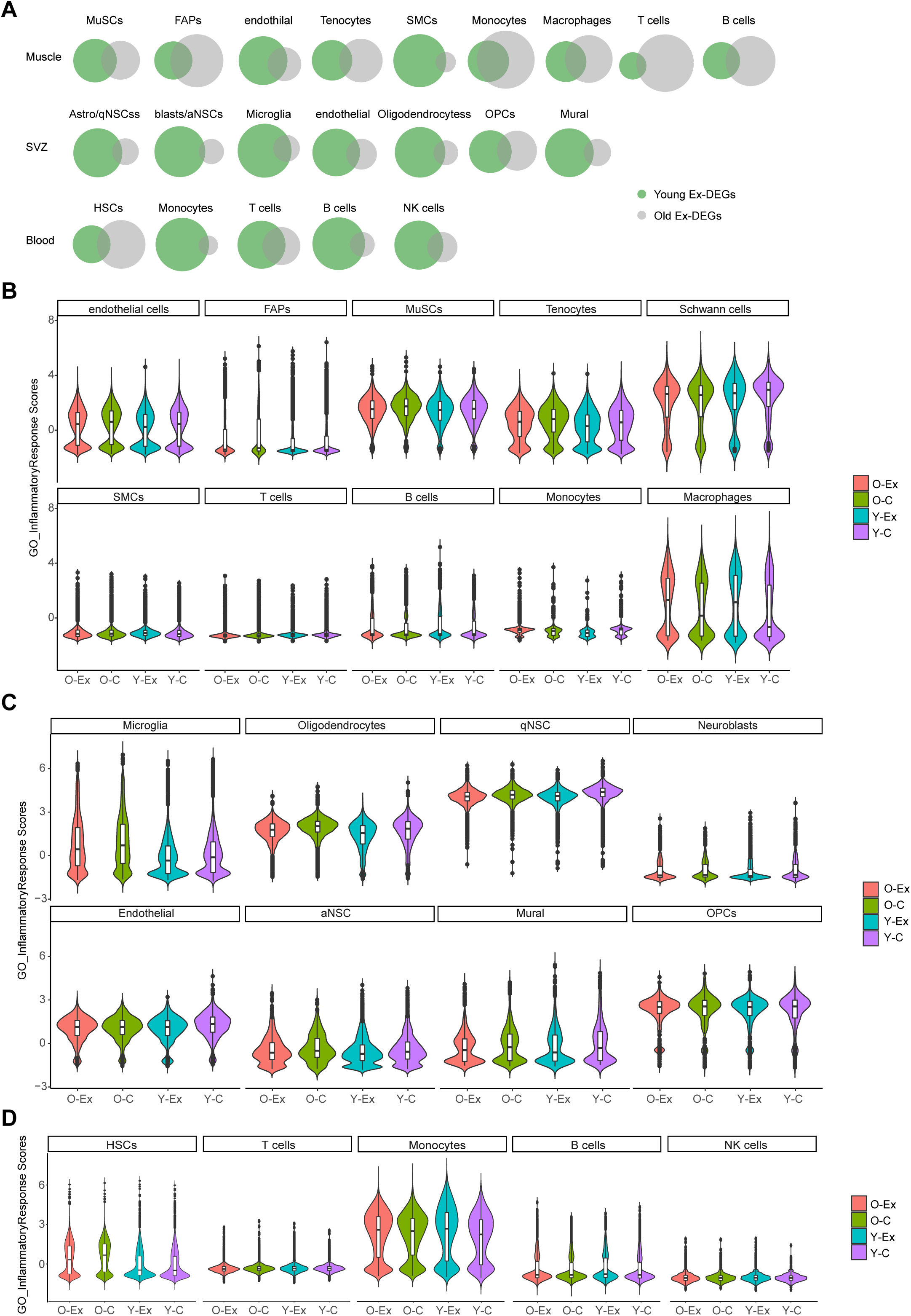
The effect of exercise on gene expression changes in various cell types. (**A**) Venn diagrams demonstrating the percentage of common Ex-DEGs of each cell type in young and old cells. **(B-D)** Violin plots showing the inflammatory scores of cells from muscle, cells from the SVZ, and hematopoietic cells.

**Figure S4:**
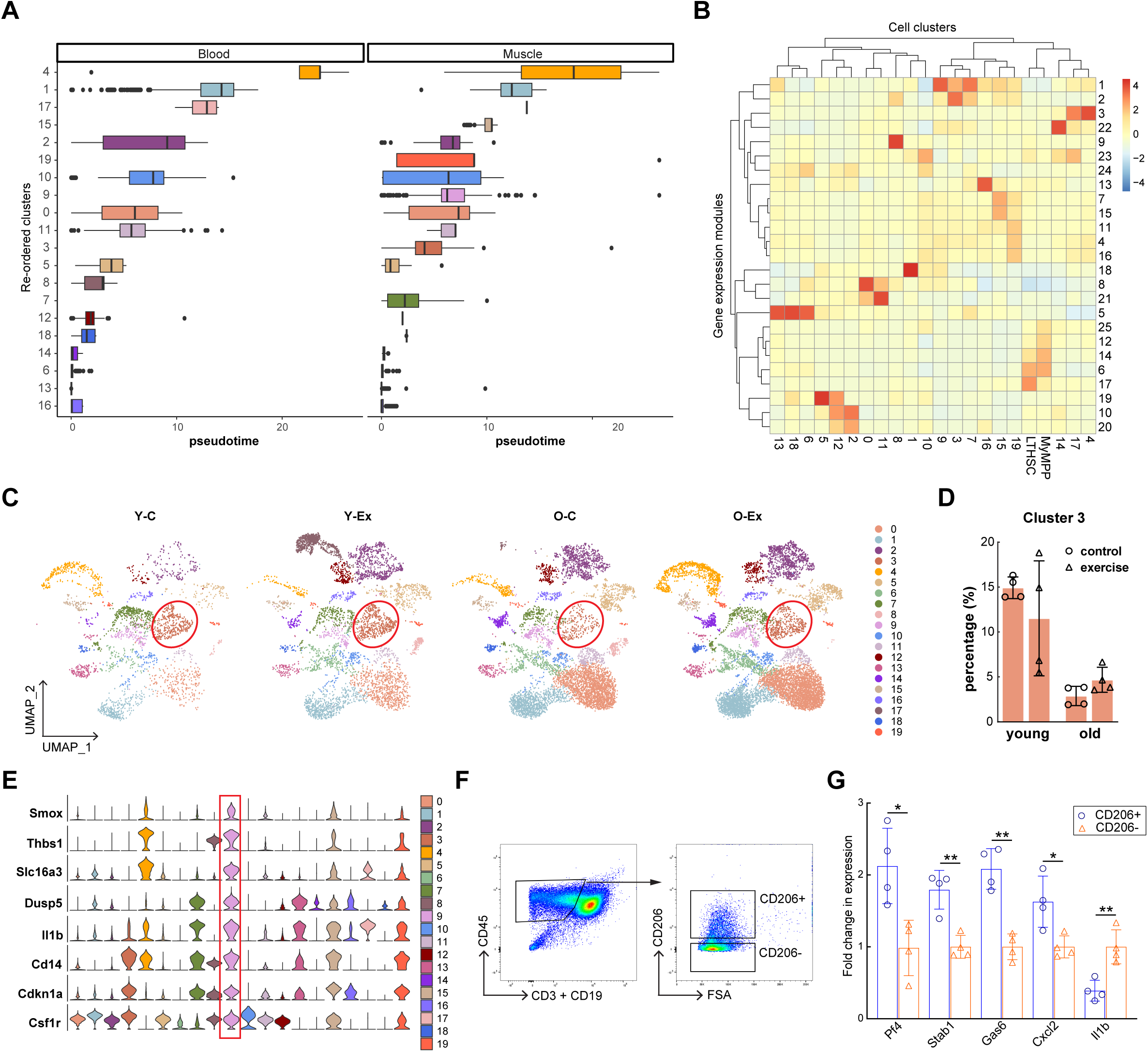
Aging and exercise-induced changes in monocytes/macrophages. (**A**) Bar graphs representing the position of each clusters in Figure 4A along the pseudotime trajectory. **(B)** Clustered heat map demonstrating the gene expression modules in all clusters in Figure 4A. **(C)** UMAPs of the 20 monocyte/macrophage clusters in Y-C, Y-Ex, O-C and O-Ex mice. Cluster 3 is marked by the red circle. **(D)** Bar graph demonstrating the relative ratio of cluster 3 in all monocytes/macrophages in Y-C, Y-Ex, O-C and O-Ex mice. (**E**) Violin plots of selected marker genes of cluster 9 of monocytes/macrophages. **(F)** Representative of FACS plots of CD45^+^/CD3^-^/CD19^-^/CD206^+^ and CD45^+^/CD3^-^/CD19^-^/CD206^-^ cells from skeletal muscle. **(G)** Bar graph demonstrating the relative expression Pf4, Stab1, Gas6, Cxcl2, and Il1b in CD206^+^ and CD206^-^ monocytes/macrophages in skeletal muscle determined by RT-qPCR analysis.

**Figure S5:**
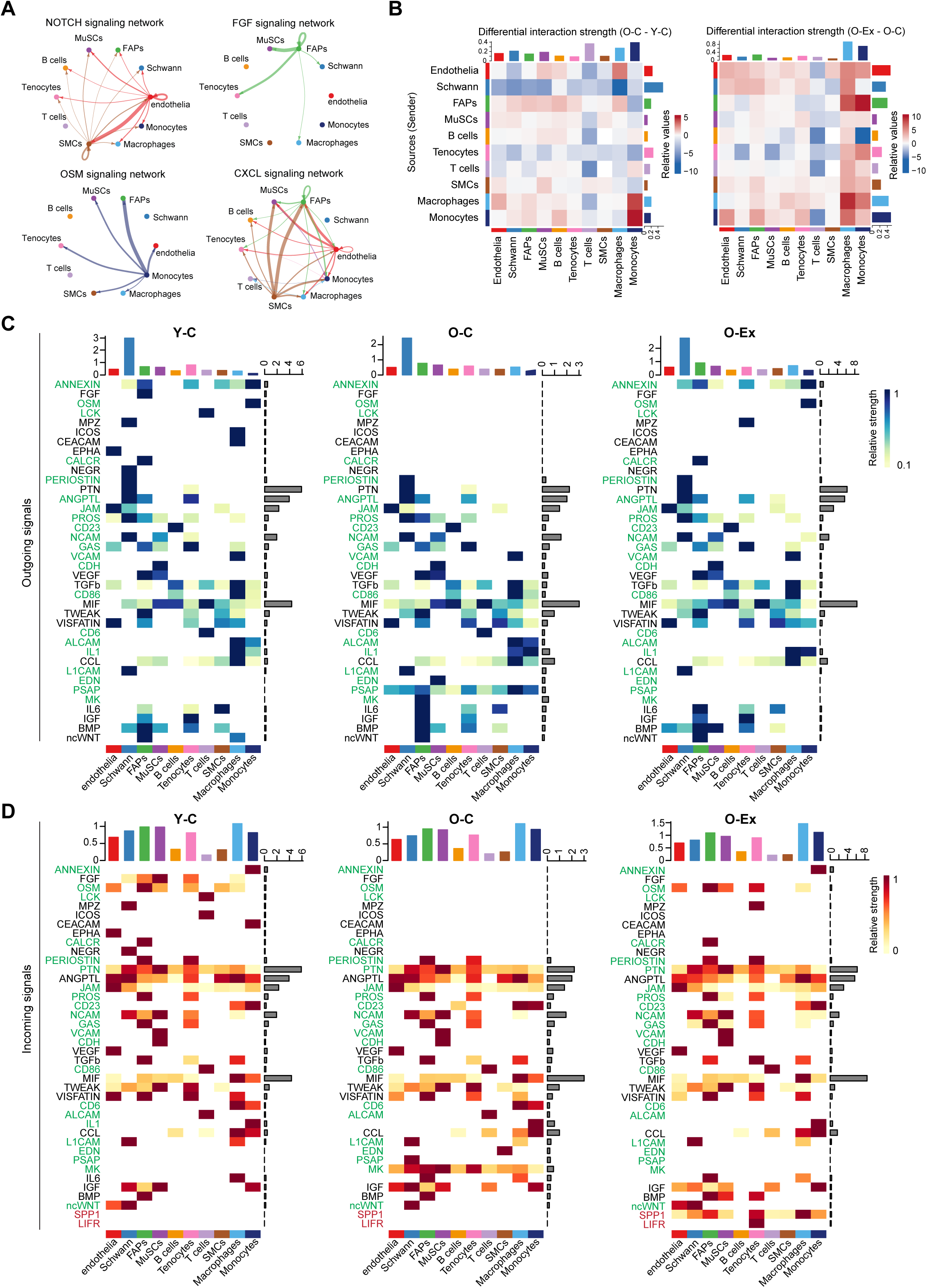
Changes in intercellular communication networks in the MuSC niche during aging and in response to exercise. (**A**) Circle plots showing the NOTCH, FGF, OSM, and CXCL signaling networks in cells from muscle of young mice. The arrows point to the cell types that express the receptors. The colors of the lines represent the source of the ligands. The thickness of the lines represents signaling strength between the signal sending and receiving cell. **(B)** Heatmaps showing the differential overall signaling strength between young and old mice (left) and between old mice without and with exercise (right). The top bars and right bars represent the sum of incoming and outgoing signaling strength of each cell type, respectively. In the left, red and blue represent higher and lower signaling strength in the O-C mice, respectively, in comparison to Y-C mice. In the right, red and blue represent lower and higher signaling strength in the O-C mice, respectively, in comparison to O-Ex mice. **(C)** Heatmaps showing the outgoing and incoming signaling pathways in various cell types in the skeletal muscle that change with age in Y-C, O-C and O-Ex mice. The pathways are ranked by the differential overall signaling flow between the Y-C and O-C conditions. The names of the pathways that were restored by exercise in old mice are labeled in green. The names of the pathways that were activated by exercise are labeled in red. The top colored bars represent the overall signaling strength in each cell type. The horizontal gray bars represent the summarized strength of each signaling pathway from all cell types in the muscle. The color scale represents the relative contribution of a cell type to the pathway.

**Figure S6:**
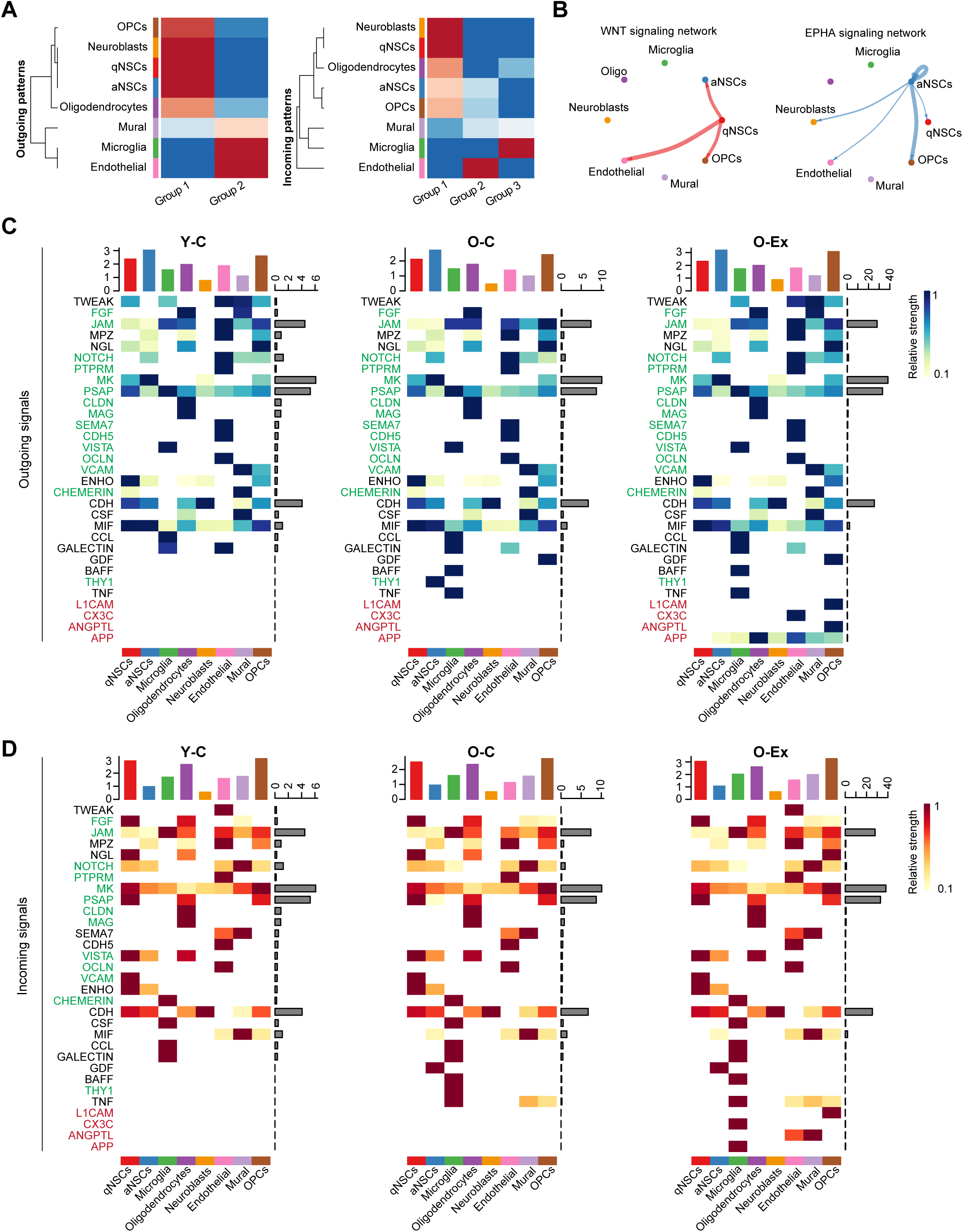
Changes in intercellular communication networks in the NSC niche during aging and in response to exercise. (**A**) Heatmaps showing the outgoing and incoming signaling patterns mediated by secreted and cell surface molecules in the SVZ. **(B)** Circle plot showing the WNT and EPHA signaling networks in the muscle of young mice. **(C)** Heatmaps showing the outgoing and incoming signaling pathways in various cell types in the SVZ that change with age in Y-C O-C and O-Ex mice. The pathways are ranked by the differential overall signaling flow between the Y-C and O-C conditions. The names of the pathways that were restored by exercise in old mice are labeled in green. The names of the pathways that were activated by exercise are labeled in red. The top colored bars represent the overall signaling strength in each cell type. The horizontal gray bars represent the summarized strength of each signaling pathway from all cell types in the muscle.

**Figure S7:**
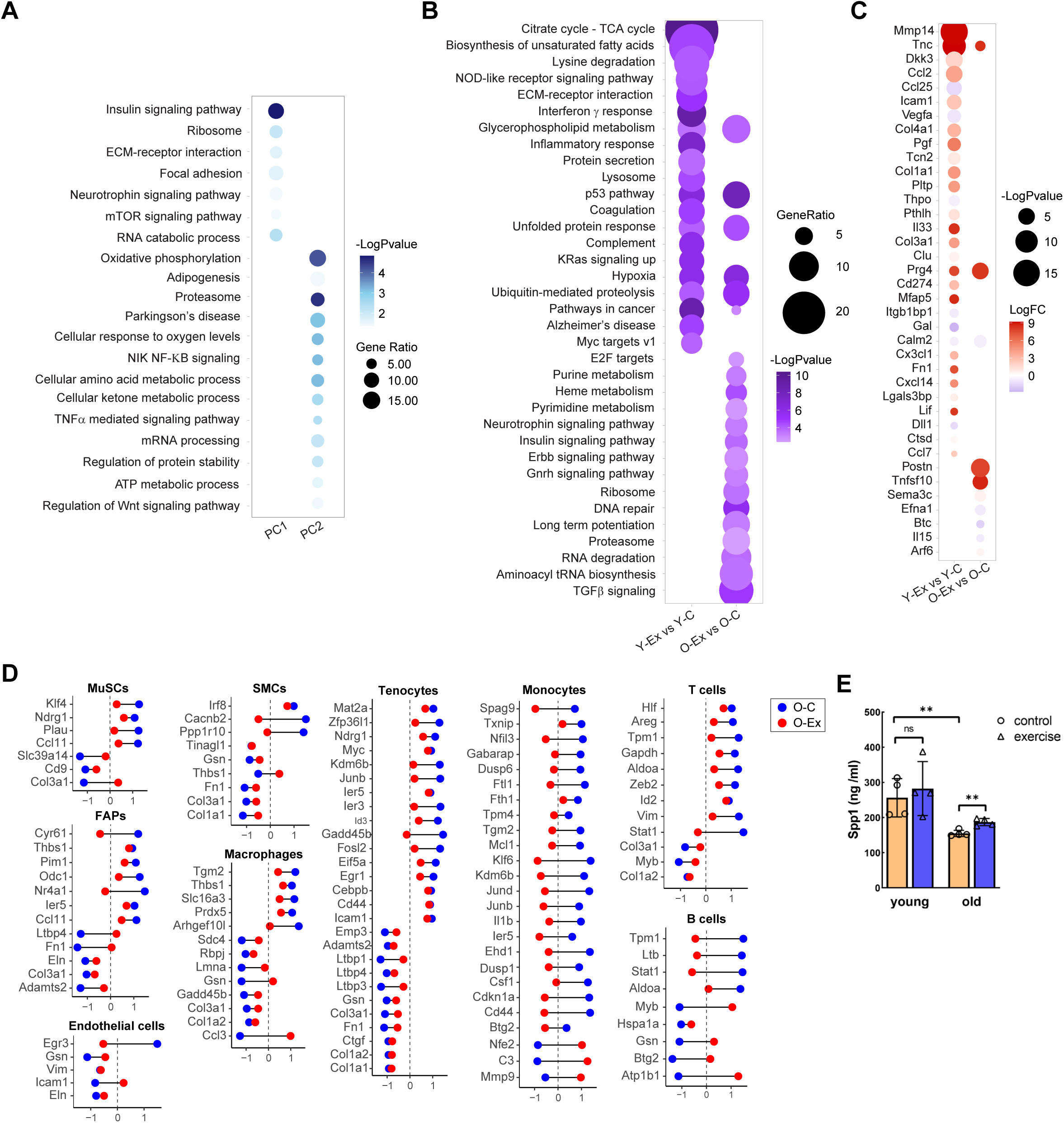
Gene expression changes in myofibers during aging and in response to exercise. (**A**) Dot plot summarizing biological pathways enriched in the two major principle components from the PCA plot in Figure 7A that distinguish fiber transcriptomes by age and exercise status. **(B)** Dot plot summarizing biological pathways enriched in exercised-induced genes in muscle fibers from young (left) and old (right) animals. **(C)** Dot plot summarizing genes encoding secreted ligands that were induced by exercise in muscle fibers from young (left) and old (right) animals. Red indicates higher expression in exercised animals. (**D**) Dumbbell plots demonstrating SPP1 target genes whose expression was reversed in muscle cell types by exercise in old mice. The x axis represents scaled relative expression to that in Y-C mice. (**E**) Bar graph showing the level of Spp1, detected by ELISA, in plasma from control and exercised young and old mice. Data are shown as mean ± SEM. *p < 0.05, **p<0.01, ***p < 0.001 (unpaired t tests).

